# The mobility of the cap domain is essential for the substrate promiscuity of a family IV esterase from sorghum rhizosphere microbiome

**DOI:** 10.1101/2022.09.10.507389

**Authors:** Marco Distaso, Isabel Cea-Rama, Cristina Coscolín, Tatyana N. Chernikova, Hai Tran, Manuel Ferrer, Julia Sanz-Aparicio, Peter N. Golyshin

## Abstract

Metagenomics offers the possibility to screen for versatile biocatalysts. In this study, the microbial community of the *Sorghum bicolor* rhizosphere was spiked with technical cashew nut shell liquid, and after incubation, the eDNA was extracted and subsequently used to build a metagenomic library. We report the biochemical features and crystal structure of a novel esterase from the family IV, EH_0_, retrieved from an uncultured sphingomonad after a functional screen in tributyrin agar plates. EH_0_ (T_opt_, 50 °C; T_m_, 55.7 °C; pH_opt_, 9.5) was stable in the presence of 10-20% v/v organic solvents and exhibited hydrolytic activity against *p*-nitrophenyl esters from acetate to palmitate, preferably butyrate (496 U mg^−1^), and a large battery of 69 structurally different esters (up to 30.2 U mg^−1^), including bis(2-hydroxyethyl)-terephthalate (0.16 ± 0.06 U mg^−1^). This broad substrate specificity contrasts with the fact that EH_0_ showed a long and narrow catalytic tunnel, whose access appears to be hindered by a thigth folding of its cap domain. We propose that this cap domain is a highly flexible structure whose opening is mediated by unique structural elements, one of which is the presence of two contiguous proline residues likely acting as possible hinges, that altogether allow for the entrance of the substrates. Therefore, this work provides a new role for the cap domain, which until now was thought to be immobile elements that contain hydrophobic patches involved in substrate pre-recognition and in turn substrate specificity within family IV esterases.

**IMPORTANCE:** A better understanding of structure–function relationships of enzymes allows revealing key structural motifs or elements. Here, we studied the structural basis of the substrate promiscuity of EH_0_, a family IV esterase, isolated from a sample of the *Sorghum bicolor* rhizosphere microbiome exposed to technical cashew nut shell liquid. The analysis of EH_0_ revealed the potential of the sorghum rhizosphere microbiome as a source of enzymes with interesting properties, such as pH and solvent tolerance and remarkably broad substrate promiscuity. Its structure resembled those of homologous proteins from mesophilic *Parvibaculum* and *Erythrobacter* spp. and hyperthermophilic *Pyrobaculum* and *Sulfolobus* spp. and had a very narrow, single-entry access tunnel to the active site, access which is controlled by a capping domain that includes a number of not conserved proline residues. These structural markers, distinct from those of other substrate promiscuous esterases, can help tuning substrate profiles beyond tunnel and active site engineering.

---

Currently, in the biotech industrial sector, there is a constant demand for novel biocatalysts with enhanced biochemical properties fulfilling industrial requirements to replace low-performance enzymes and dismiss chemical processes (1). From this perspective, the enzymatic diversity within the microbial communities, that may reach for example up to 10^3^-10^5^ microbial species in 1 g of soil, represents a yet unexplored and exceptional resource for new biocatalysts (2, 3), and thus, it can be a source of biocatalysts. A straightforward evaluation of such diversity can be performed by metagenomics, through a number of bioinformatics and so-called naïve screening methods (2, 4–6).

Carboxylic ester hydrolases (EC 3.1.1.-) are a class of enzymes widely distributed in nature that have attracted much attention because of their high sequence diversity, structural versatility, catalytic versatility, robustness and lack of need for cofactors (7–10). Considering the importance of these classes of enzymes, it is not surprising that they are priority targets to screen by metagenomics. For example, approximately 280,638 sequences encoding carboxylic ester hydrolases have been cataloged (10), and more than 4,000 new enzymes have been discovered through metagenomics (11); few of them show enhanced biophysical properties, such as stability at extreme temperatures, salinity, pH and solvents conditions, high catalytic efficiency, broad substrate specificity and regio- and enantioselectivity (6, 12–14). Additionally, metagenomic studies on soil environments significantly contributed to broadening the knowledge about this class of enzymes with the reannotation of sequences known as hypothetical proteins and the definition of new esterase families (15, 16).

Carboxylic ester hydrolases are also the most studied enzymes with respect to promiscuity (17), including catalytic (18) and substrate (19) promiscuity. The question that arises is, would it not be more efficient to have just a couple of promiscuous enzymes with the ability to perform a wide range of reactions (19). Based on this principle, biotechnological industries demand biocatalysts that are capable of transforming a wide range of substrates. Their isolation has been solved through the application of metagenomic techniques that have emerged as robust tools for the discovery of unknown biocatalysts, including ester hydrolases with a broad substrate scope. This method, together with the understanding of the structural mechanisms and engineering behind promiscuous enzymes, seems crucial for their industrial implementation. In this context, our recent study of approximately 145 ester hydrolases has revealed that members of family IV show higher levels of substrate promiscuity than those of other families. This is the result of their capacity to adapt the topology of the large but occluded active sites to a high variety of substrates but also to the presence of a cap domain next to the entrance of the substrate tunnel (19). The cap domains were thought to be immobile elements, unlike lid elements in lipases, that contain hydrophobic patches involved in substrate pre-recognition which could contribute to the substrate promiscuity within members of family IV (20).

The aim of this study was to mine carboxylic ester hydrolases of the soil rhizosphere of sorghum plants after enrichment with cashew nut shell liquid (CNSL), an oily substance rich in phenolics and fatty acids (21), with respect to novel biocatalysts. By applying an activity-based strategy, four such enzymes were retrieved and characterized. We also reported the crystal structure of one of these enzymes with an extraordinarily broad substrate specificity. The structural information, complemented with the experimental analysis of the substrate specificity of a series of variants, provides additional mechanistic understanding of the molecular basis for the role of the cap domain in the promiscuous character of members comprising family IV esterases. Importantly, we propose the role of proline residues at the N-terminal region comprising α1-α2 helices as hinges of the movement of the cap domain allowing substrates to enter the active center.

## Results

### Identification of a carboxylic ester hydrolase

The search for hydrolytic enzymes in the metagenomics library SorRhizCNSL3 W (see details in Materials and Methods) was carried out by applying a metagenomic functional screening approach. The strategy involved the use of LB agar plates containing tributyrin (22). The pooling of 40 clones per well dispensed into 96-well microtiter plates (approximately 3.8 k clones in one plate) facilitated the colony screening at a high throughput. After screening more than 100,000 clones, 1 hit from the SorRhizCNSL3 W library showed lipase/esterase activity. Fosmid insert was extracted using the QUIAGEN Large-Construct Kit (QIAGEN, Hilden, Germany) according to the manufacturer’s protocol, digested with restriction enzymes for insert size estimation, and the insert sequenced by Illumina technology. Upon the completion of sequencing, the reads were quality-filtered and assembled to generate nonredundant meta-sequences, and genes were predicted and annotated as described previously (12). One gene encoding a predicted carboxylic ester hydrolase was identified. The gene was amplified with specific primers, cloned into the p15TV-L vector and transformed into *E. coli* BL21(DE3) cells for expression of the N-terminal (His)6-tagged proteins. The deduced amino acid sequence of the enzyme (324 amino acids long) was used for homology searches in the taxonomy and functional assignment. A database search indicated that EH_0_ showed 99% identity with the α/β hydrolase enzyme from *Sphingomonas pruni* (protein ID WP_066587239), both of which are classified as members of the α/β hydrolase-3 family (PF07859). The typical H-G-G-G motif and the G-X-S-X-G catalytic motif are conserved, and EH_0_ clustered together with family IV esterase.

### Biochemical characterization

The recombinant protein was successfully expressed in soluble form and purified by nickel affinity chromatography. Purified protein was desalted by ultrafiltration, and its enzymatic activity was assessed. Four model *p*-nitrophenyl (*p-*NP) ester substrates with different chain lengths were used to determine the substrate specificity of the enzymes and therefore determine whether the enzymes are in fact true lipases or esterases. Lipases hydrolyze ester bonds of long-chain triglycerides more efficiently than esterases, which instead exhibit the highest activity toward water soluble esters with short fatty acid chains (23). The substrates used for the hydrolytic test were *p-*NP acetate (C2), *p-* NP butyrate (C4), *p-*NP dodecanoate (C12), and *p-*NP palmitate (C16). The hydrolytic activity was recorded under standard assay conditions (Fig. 1). EH_0_ showed a specific activity of 496.5 U mg^−1^ for *p*-NP butyrate, which was the best substrate. Lower levels of activity were observed with longer chain esters (C ≥ 12). The esterase followed the Michaelis–Menten kinetics, and its kinetic parameters are reflected in Table 1. A comparison of the catalytic efficiency values (*k_cat_*/K_M_) indicated a high reactivity toward *p*-NP-butyrate followed by *p*-NP-acetate.

**FIG 1.**
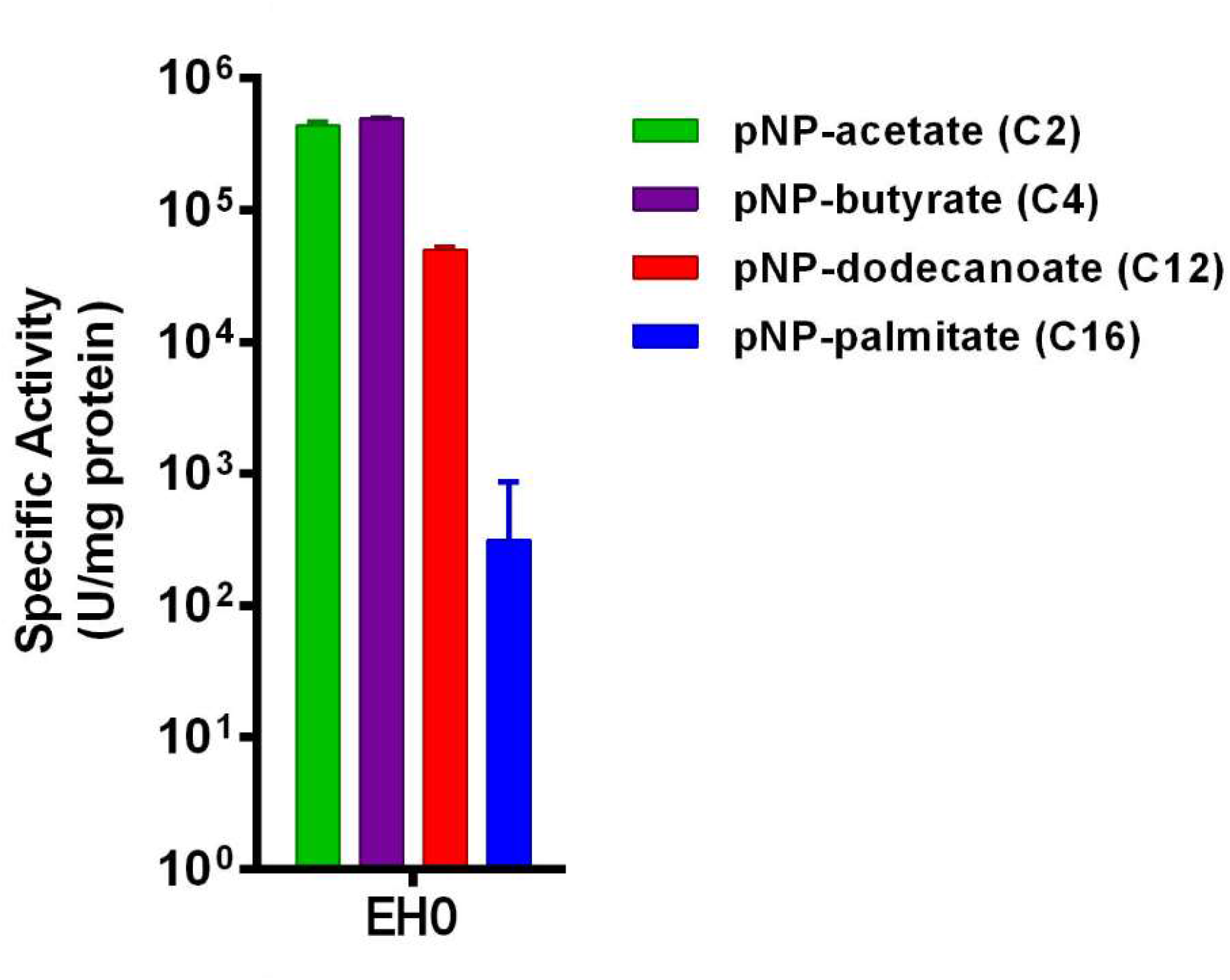
Substrate selectivity of EH_0_ against a set of model *p*-NP esters of different acyl chain lengths. Specific activities (mean ± SD of triplicates) are shown. Reactions contained 1 mM of the corresponding *p*-NP esters and were conducted in the presence of 1-5% DMSO/acetonitrile (see Methods), under standard conditions described in Materials and Methods. At the solvent concentration used the enzyme 100% of its activity compared to a control without solvent (Table S2).

**TABLE 1.** Kinetic parameters of EH_0_ against *p*NP-esters. The results are the mean ± SD of triplicates.

| Substrate<sup>1</sup> | K<sub>M</sub> (mM) | V<sub>max</sub> (μM min<sup>-1</sup>) | k<sub>cat</sub> (min<sup>-1</sup>) | k<sub>cat</sub>/K<sub>M</sub> (s<sup>-1</sup> mM<sup>-1</sup>) |
| --- | --- | --- | --- | --- |
| *p*-NP acetate | 1.61 ± 0.12 | 291.50 ± 7.62 | 41.64 ± 1.09 | 0.430 |
| *p*-NP butyrate | 1.08 ± 0.18 | 252.60 ± 13.84 | 36.09 ± 1.98 | 0.558 |
| *p*-NP dodecanoate | 1.00 ± 0.28 | 25.56 ± 1.80 | 3.65 ± 0.26 | 0.061 |

<sup>1</sup>Assays were performed in the presence of 5.0-7.5% DMSO/acetonitrile (see Materials and Methods), concentration at which the enzyme 100% of its activity compared to a control without solvent (Table S2).

Its voluminous (volume of the active site cavity: 5133 Å^3^) but low exposed (solvent accessible surface area (SASA): 5.07 over 100 dimensionless percentage) active site allows hydrolysis of a broad range of 68 out of 96 structurally and chemically diverse esters (Table S1), as determined by a pH indicator assay (pH 8.0, 30 °C). Phenyl acetate (30.23 U mg^−1^) and glyceryl tripropionate (29.43 U mg^−1^), were the best substrates. We also found that EH_0_ efficiently hydrolyzed bis(2-hydroxyethyl)-terephthalate (BHET; 163.6 ± 6.2 U mg^−1^), an intermediate in the degradation of polyethylene terephthalate (PET) (24); HPLC analysis (Fig. S1), performed as described (25), confirmed the hydrolysis of BHET to mono-(2-hydroxyethyl)-terephthalic acid (MHET) but not to terephthalic acid (TA). However, using previously described conditions (25), we found that the enzyme did not hydrolyze large plastic materials such as amorphous and crystalline PET film and PET nanoparticles from Goodfellow. According to the number of hydrolyzed esters, EH_0_ can be thus considered as an esterase with a wide substrate specificity, similar to other enzymes of the family IV (19, 20).

EH_0_ showed maximal activity at 50 °C, retaining more than 80% of the maximum activity at 40-55 °C (Fig. 2A), suggesting that it is moderately thermostable. This was confirmed by circular dichroism analysis, which revealed a denaturing temperature of 55.7 ± 0.2 °C (Fig. 2B). Its optimal pH for activity was 9.5 (Fig. 2C). The effect on the enzymatic activity of organic solvents at different concentrations was evaluated (Table S2). An activation effect was observed for EH_0_ when 10% methanol (60% activity increase) and 10-20% DMSO (22-40% increase) were added to the reaction mixture. The presence of bivalent and trivalent cations did not have a remarkable positive effect on the activity of the enzymes, which showed, in some cases, tolerance to high concentrations of cations (Table S3). A prominent inhibiting effect was shown for all cations, except for magnesium, which was well tolerated at 1-10 mM (<5% inhibition).

**FIG 2.**
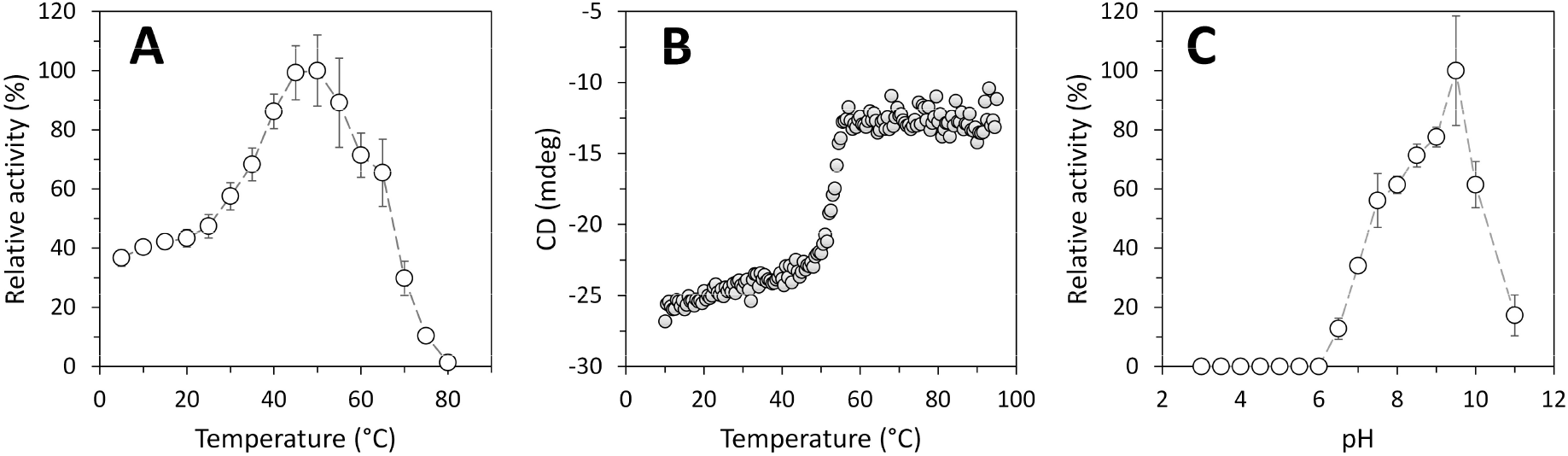
Optimal parameters for the activity and stability of purified EH_0_. (A) Temperature profile. (B) The thermal denaturation curve of EH_0_ at pH 7.0 was measured by ellipticity changes at 220 nm and obtained at different temperatures. (C) The pH profile. The maximal activity was defined as 100%, and the relative activity is shown as the percentage of maximal activity (mean ± SD of triplicates) determined under standard reaction conditions with *p*-NP butyrate as the substrate. Graphics were created with SigmaPlot version 14.0 (the data were not fitted to any model).

### EH_0_ presents tight folding of its cap domain

The crystal structure of wild type EH_0_ was obtained at 2.01 Å resolution, with the P2_1_2_1_2_1_ space group and two crystallography-independent molecules in the asymmetric unit. Molecular replacement was performed using Est8 as a template (PDB code 4YPV) (26), and the final model was refined to a crystallographic R-factor of 0.1717 and R-free of 0.2019 (Table S4). As with other reported family IV esterases, EH_0_ has an **α**/**β** hydrolase fold with two different domains, a cap domain (residues 1-43 and 208-229) and a catalytic domain (residues 44-207 and 230-324), constituted by a total of 9 **α**-helices and 8 **β**-sheets (Fig. 3A). The catalytic domain was composed of a central **β**-sheet with eight parallel **β**-strands (**β**1, **β**3, **β**4, **β**5, **β**6, **β**7, **β**8), except **β**2, which was antiparallel and surrounded by five **α**-helices (**α**3, **α**4, **α**5, **α**8, **α**9). The cap domain involved four **α** helices (**α**1, **α**2, **α**6, **α**7) (Fig. 3B). There were two *cis* peptides, Ala122-Pro123 and Trp127-Pro128, located at the **β**4-**α**4 turn within the catalytic domain.

**FIG 3.**
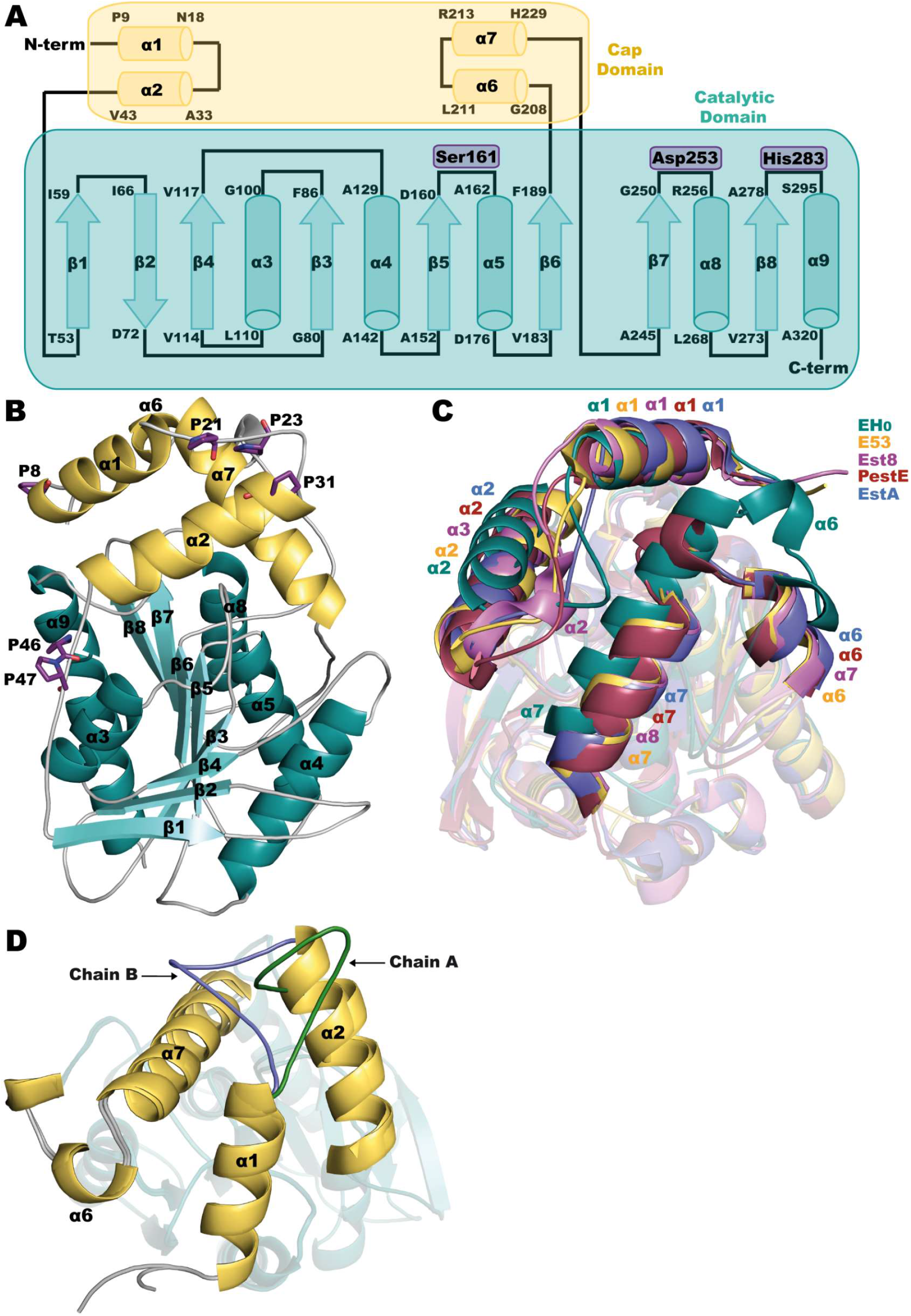
A) Topology diagram of EH_0_. The catalytic triad is highlighted in violet. B) EH_0_ folding, with the cap domain in yellow and the catalytic domain in teal. The prolines within the N-terminal cap domain are depicted as violet sticks. C) Superposition of EH_0_ and its homologs: E53 (yellow, PDB code 7W8N), Est8 (violet, PDB code 4YPV), PestE (raspberry, PDB code 3ZWQ) and EstA (slate, PDB code 5LK6). D) Superposition of subunits A and B showing different conformations of loop α1-α2, with chain A in green and chain B in blue. Cap/catalytic domains follow the color code described in B.

Analysis of EH_0_ folding using the *DALI* server (27) was employed to search for homologous proteins. The closest homologs are E53 isolated from *Erythrobacter longus*, with 46% identity and RMSD of 2.3 Å on 296 Cα atoms (PDB code 7W8N) (28); Est8 isolated from *Parvibaculum*, with 38% identity and RMSD of 2.5Å on 291 Cα atoms (PDB code 4YPV) (26); PestE isolated from *Pyrobaculum calidifontis*, with 33% identity and RMSD of 2.6Å on 294 Cα atoms (PDB code 3ZWQ) (29); and EstA isolated from *Sulfolobus islandicus REY15A*, with 31% identity and RMSD of 2.6Å on 281 Cα atoms (PDB code 5LK6). The structural superimposition of these proteins reveals a high conservation of the corresponding catalytic domains, and also a common spatial arrangement of the helices at the cap domains in all the proteins except EH_0_, where α2 and the long α7 are visibly shifted very close to its EH_0_ active site and apparently impeding the entrance of substrates (Fig. 3C). This was unexpected considering the broad substrate specificity of esterase EH_0_, approaching that of most promiscuous ones (19). Thus, withdrawal of the cap domain seems a necessary requirement for allowing access of the bulky substrates to the EH_0_ catalytic site.

In agreement with this assumption, a substantial rearrangement of the cap domain was previously described in the homolog esterase EST2 (having 31 % sequence identity), with its M211S/R215L double variant being trapped in the crystal in a conformation resembling the open form of lipases (30). However, the authors did not assign any biological relevance to this issue, considering this state as an artifact derived from the crystal packing. In our study, some flexibility at this region has been found from the two conformations adopted by loop α1-α2 observed in subunits A and B of EH_0_ (Fig. 3D). In addition, the inspection of residues within the cap domain showed a high number of proline residues (Pro8, Pro21, Pro23, Pro31, Pro46 and Pro47) (Fig. 3B) that are mostly non-conserved and could confer flexibility to the N-terminal cap domain allowing substrate entrance. This feature will be discussed below.

### EH_0_ is a dimeric enzyme

While E53 and Est8 are monomeric enzymes, and EstA is a tetramer, EH_0_ is presented as a biological homodimer with approximate dimensions of 6.3 × 4.9 × 2.7 nm, which is assembled in a twofold axis symmetry arrangement (Fig. 4A) that buries 5.6% of its total surface area. Hydrogen bonds mainly involve β8 and α8, while salt bridges involve motifs β8 and α9 (Fig. 4B and Table S5). Similar to its dimeric homolog PestE, oligomerization occurs through a tight interaction of β8 strands from both subunits. However, while only the subsequent helices **α**12 were involved in the PestE interface (29) (Fig. 4C), both the precedent **α**8 and the subsequent **α**9 helices make the EH_0_ interface (Fig. 4D). These different interactions observed in PestE and EH_0_ were reflected in a different orientation between the monomers that, nevertheless, present a similar distance between the catalytic serines of 35-38 Å and an equivalent disposition of the tunnels giving access to the catalytic site at two edges of the dimer (Fig. 4E, 4F). Therefore, as seen in Fig. 4A, the two cap domains are far from the interface and project out from the dimer, revealing that dimerization is not affecting the cap function.

**FIG 4.**
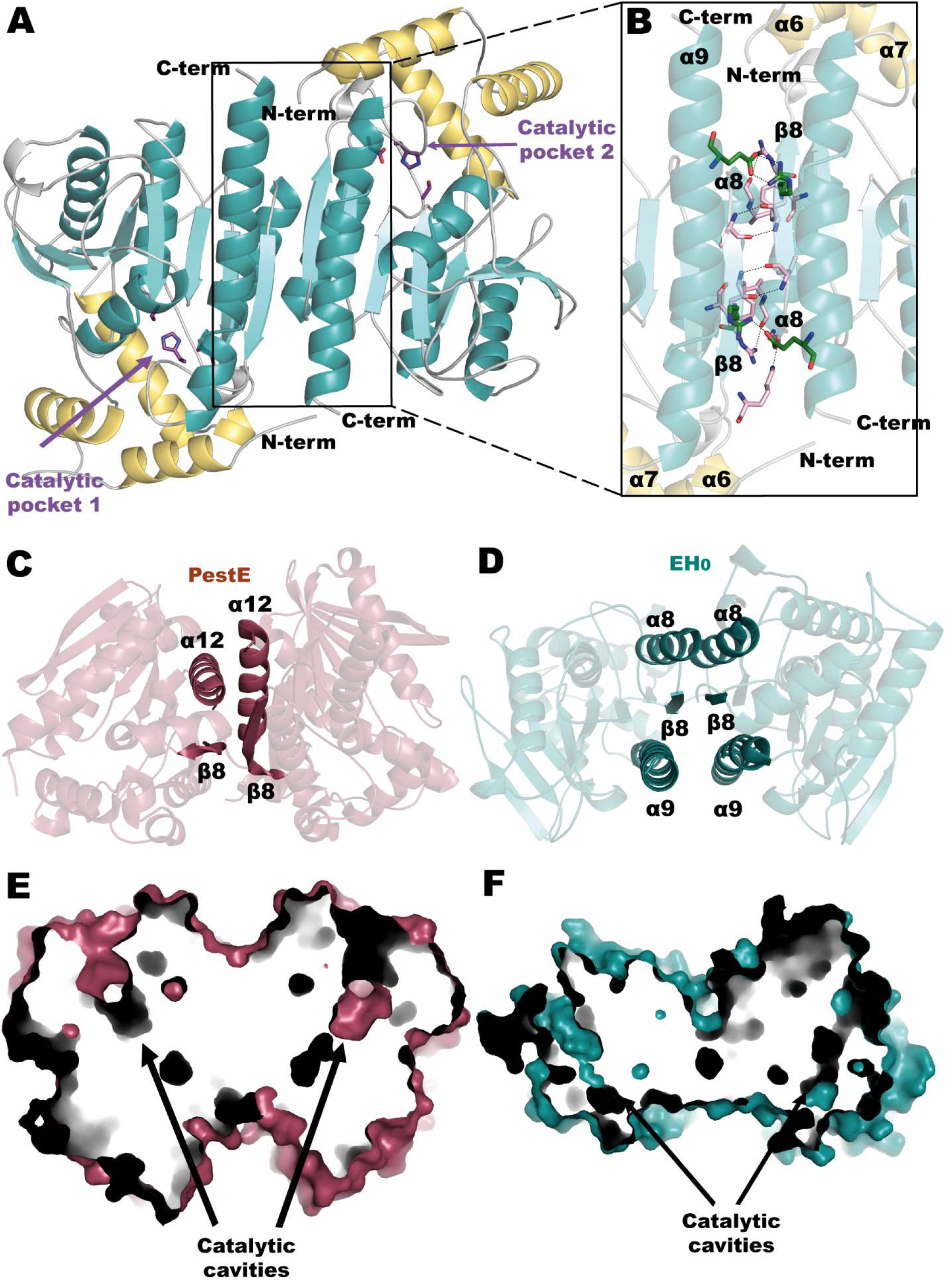
A) EH_0_ homodimer showing the catalytic triad as sticks (violet). Both subunits are related by a twofold axis that is perpendicular to the view shown and situates the two active sites on opposite faces of the dimer. B) A magnified image of the dimeric interface that covers 722.9Å^2^ (approximately 5.6% of the total surface), showing hydrogen bonds (pink) and salt bridges (green). The Cap domain is colored yellow, whereas the catalytic domain is shown in teal. C) Dimeric interface of the PestE homolog where the interactions are made by β8 and α12. D) Dimeric interface of EH_0_, where the interactions are made by α8, β8 and α9. Disposition of catalytic tunnels from both subunits in PestE (E) and EH_0_ (F).

### The peculiar EH_0_ active site

EH_0_ has a long catalytic tunnel with a very narrow entrance (approximately 16.8 Å depth) (Fig. 5A). The catalytic triad of EH_0_ is formed by Ser161 (in the conserved motif ^159^G-D-S-A-G^163^ in the nucleophilic elbow), Asp253 (in the conserved motif ^253^D-P-I-R-D^258^) and His283 (Fig. 5B). To analyze the active site, a series of soaking and cocrystallization experiments were performed with different suicide inhibitors, all of which were unsuccessful. A deep inspection of the active site showed that the nucleophilic Ser161 is hydrogen bonded to Glu226 from the close **α**7, and a movement of loop **β**3-**α**3, including the oxoanion in the conserved motif ^87^H-G-G-G^90,^ compared to EH_0_ homologs (Fig. 5B). Therefore, to remove this hydrogen link in an attempt to expand access to the active site enabling ligand binding, the EH_0E226A_ variant was produced by directed mutagenesis. However, the crystals grown from this variant failed to diffract, suggesting a high level of disorder, putatively due to an increased mobility of the cap. Interestingly, the activity profile of this variant reveals a faster hydrolysis of most of the esters (62 esters in total; from 70- to 1.2-fold; average: 6.1-fold). Only 7 esters were hydrolyzed to a lower extent compared to the wild type (from 1.5- to 17-fold; average: 4.6-fold). Remarkably, the EH_0E226A_ variant was capable of accepting large and voluminous esters such as dodecanoyl acetate, pentadecyl acetate, vinyl laurate, methyl 2,5-dihydroxycinnamate, and ethyl 2-chlorobenzoate, which were not accepted by the wild-type enzyme (Table S1). This apparently supports that removal of the Ser161-Glu226 hydrogen bond increases cap flexibility and enhances the enzymatic efficiency toward large esters either at the acyl or alcohol sites.

**FIG 5.**
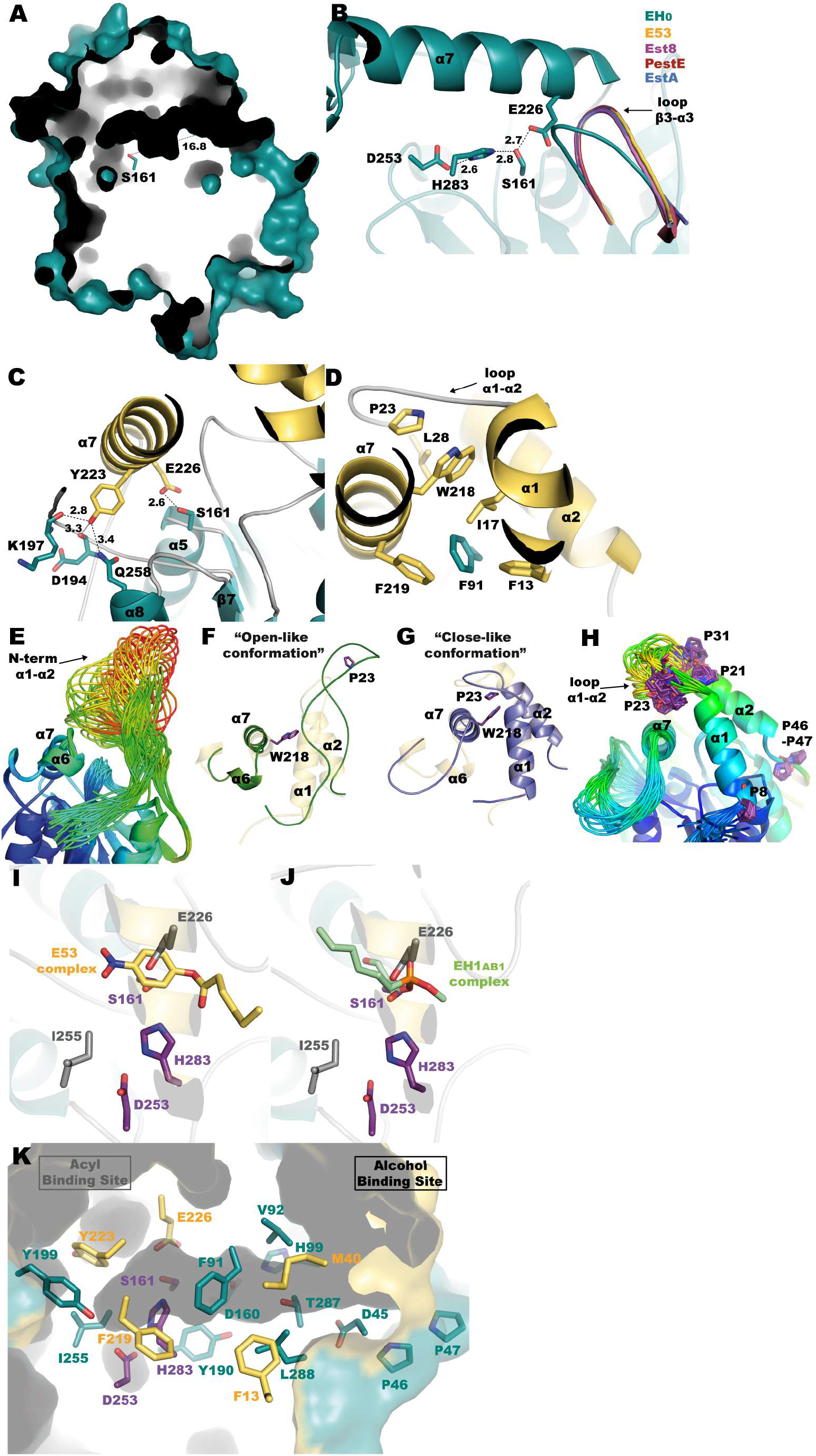
A) A tunnel 16.8 deep giving access to the catalytic triad. B) Hydrogen bond network among the residues making the catalytic triad of EH_0_ (teal). Movement of loop β3-α3 in EH_0_ (housing the oxoanion) with respect to its homologs E53 complexed with (4-nitrophenyl)hexanoate (yellow), Est8 (violet), PestE (raspberry) and EstA (slate). C) Hydrogen bond network of residues Glu226 and Tyr223 from α7. D) Hydrophobic patch where Trp218 is located. The EH_0_ cap domain is shown in yellow and the catalytic domain in teal (C and D.)E) All possible conformations of the region comprising α1-α2 shown by the ensemble refinement of molecule A. Color code gives chain mobility, from low (blue) to high (red). The opposed scenarios of the region α1-α2, from an “open conformation” (F, green) to a “closed conformation” (G, slate). The cap domain seen in the crystal structure is shown in yellow. H) All conformers shown by the ensemble refinement of molecule B. I and J) Superimposition of EH_0_ onto E53 complexed with 4-nitrophenylhexanoate (yellow) and EH_1AB1_ in complex with a derivative of methyl 4-nitrophenylhexylphosphonate (green), showing only the ligands. The catalytic triad of EH_0_ is shown as violet sticks, and residues Glu226 and Ile255 are shown as gray sticks. The EH_0_ cap domain is shown in yellow and the catalytic domain in teal. K) Active site cavity of EH_0_ with acyl and alcohol moieties. Residues from the cap domain are colored yellow, and residues from the catalytic domain are colored teal. The catalytic triad is shown in violet.

Indeed, it appears clear that helix α7 must retract from the catalytic site to allow substrate entrance, but this helix seems fixed by many atomic interactions. Close to the Ser161-Glu226 hydrogen bond, Tyr223 makes additional hydrogen bonds to Asp194 and Lys197 main chain, from loop β6-α6, and with the side chain of Gln258 from α8 (Fig. 5C). A mutation EH_0Y223A_ was generated by directed mutagenesis and found, unlike the EH_0E226A_ mutation, little effect on conversion rates compared to the wild-type enzyme. However, this mutation allowed the hydrolysis of large and voluminous esters, such as dodecanoyl acetate, pentadecyl acetate, methyl 2,5-dihydroxycinnamate and ethyl 2-chlorobenzoate, which were also hydrolyzed by the variant EH_0E226A_. Unlike EH_0E226A_, the EH_0Y223A_ variant was not unable to hydrolyze vinyl laurate, but it was able to hydrolyze methyl 3-hydroxybenzoate, not accepted by the EH_0E226A_ variant. This suggested that Tyr223 may play a role in accepting large esters, particularly at the acyl side, but may also play an additional role in substrate specificity different from that of Glu226.

Additionally, at the beginning of α7, Trp218 is within a hydrophobic pocket surrounded by residues Phe13, Ile17, Leu28 and Phe219 from the cap domain and Phe91 from the catalytic domain, which is also anchored by interaction with loop α1-α2 and Pro23 (Fig. 5D). Thus, to depict how these tight molecular packing may be disrupted by the proposed cap motion, molecular dynamics were applied to crystallographic refinement through the ensemble refinement strategy, which is shown to model the intrinsic disorder of macromolecules giving more accurate structures. The ensemble models obtained for molecules A and B within the asymmetry unit are shown in Fig. 5E and 5H, respectively. The analysis of the molecule A conformers (Fig. 5E) revealed that the region comprising α1 and α2 shows a wide spectrum of possible pathways from more “open” to more “closed” conformations. At one edge, Pro23 is in an extended α1-α2 loop far from Trp218, therefore releasing α7 that consequently could retract from the catalytic pocket (“open-like conformation”) (Fig. 5F). In fact, the ensemble refinement models a very flexible conformation being even unstructured at regions corresponding to α1 and α2. At the other edge, the second scenario is the entrapment of Trp218 by Pro23 in loop α1-α2, hindering substrate entrance (“closed-like conformation”) (Fig. 5G). This last scenario is equivalent to the 3D structure captured by crystallography (Fig. 5D). Furthermore, three prolines can be found within this α1-α2 loop, Pro21 (at the end of α1), Pro23 (at the middle) and Pro31 (at the beginning of α2), all of them unique to EH_0,_ which are probably behind the two different conformations observed at this loop in both subunits within the asymmetric unit (Fig. 3B and 3D), and explain the ensemble of conformers modelled for molecule B (Fig. 5H).

Furthermore, as seen in Fig. 3B, Pro46 and Pro47 are potential hinges that would involve flexibility of a larger region of the EH_0_ cap domain, including the whole N-terminal peptide chain up to the end of α2. This is consistent with the more “open” conformations resulting from the ensemble refinement shown in Fig. 5F. In fact, the sequence comparison of EH_0_ to its closest homologs reveals that only EH_0_ has two sequential prolines at this region and that Pro46 is unique to EH_0_ (Fig. 6), which could be a reason behind the high EH_0_ promiscuity. Interestingly, EST2 also presents the two contiguous Pro residues (Pro38, Pro39), which can also confer high mobility to its cap domain and facilitate the “open-like” conformation captured in the crystal mentioned above (30). Therefore, the mutation EH_0P46A_ was generated by directed mutagenesis and submitted to crystallization experiments to investigate the Pro46 putative role. However, the crystals grown from this variant EH_0P46A_ failed to diffract, suggesting that removal of Pro46 introduces some structural instability to the polypeptide chain resulting in crystal disorder. Moreover, analysis of the activity profile showed that Pro46 is a critical residue for the entry and hydrolysis of bulky substrates, as its mutation by Ala extends the substrate specificity from 68 to 84 esters. Additionally, the hydrolytic rate increased from 1.2- to 18000-fold (average: 335-fold) for most esters. This variant was also able to hydrolyze large glyceryl trioctanoate and 2,4-dichlorophenyl 2,4-dichlorobenzoate, which were not hydrolyzed by the wild-type enzyme or the EH_0E226A_ and EH_0Y223A_ variants. Consequently, although the proposed role of Pro46 and Pro47 as putative hinges enabling the opening of the cap domain seems appealing, other mechanisms promoting EH_0_ plasticity to bulky substrates may also operate. Furthermore, as seen below, it should be noted that the two proline residues are located at the entrance of the narrow tunnel giving access to the active-site, an issue that ascribes a prominent role to both residues in binding activity and specificity.

**FIG 6.**
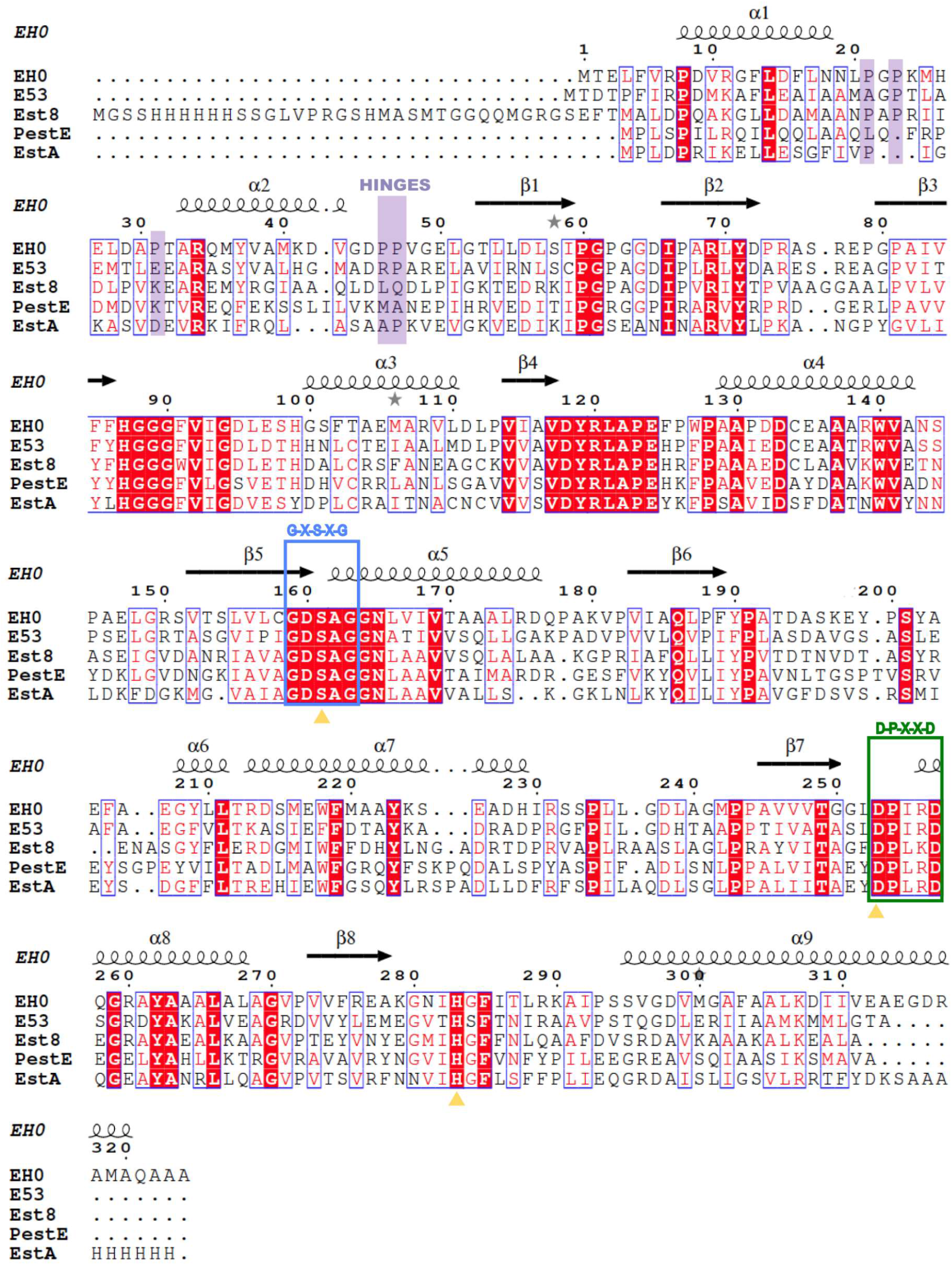
A) Multiple sequence alignment of EH_0_ and its homologs E53, Est8, PestE and EstA. Highlighted in boxes are the conserved motif G-X-S-X-G (in blue), including the catalytic serine, and the D-P-X-X-D (in green), including the catalytic aspartic acid. The prolines from loop α1-α2 (Pro21, Pro23 and Pro31) and the potential hinges (Pro46 and Pro47) are underlined in purple. The catalytic triad residues are shown with yellow triangles.

Structural details of the EH_0_ active site and assignment of the acyl and alcohol moieties were explored by comparison with its homologs E53 complexed with 4-nitrophenyl hexanoate (PDB code 6KEU) (28) and EH_1AB1_ in complex with a derivative of methyl 4-nitrophenylhexylphosphonate (PDB code 6RB0) (31). However, the acyl and alcohol moieties of these complexes are located in opposite sites (Fig. 5I, 5J). Therefore, as we could not obtain complexes from EH_0_, the activity experiments were crucial to correctly assign acyl/alcohol moieties. As mentioned above, experimental evidence demonstrated that Tyr223 produces a steric hindrance at the acyl moiety, and consequently, the acyl and alcohol sites correspond to those observed in EH_1AB1_ (Fig. 5J). On the basis of this assumption, the acyl binding site seems to be a small cavity bordered by the Tyr199 and Ile255 side chains, which produce steric hindrance for substrates with large acyl moieties. The long and narrow alcohol binding site is surrounded by both hydrophobic (Phe13, Phe91, Val92, Leu288) and hydrophilic residues (Asp45, His99, Asp160, Tyr190, Thr287), with Pro46 and Pro47 being at the entrance of the tunnel (Fig. 5K). Most residues at the acyl and alcohol moieties are conserved among EH_0_ homologs, with the exception of Met40, Pro46, Tyr199, Glu226 and Thr287. Remarkably, the bulky Met40 and Tyr199 residues are substituted by smaller residues in the EH_0_ homologs. As previously mentioned, Pro46 is unique in EH_0_, while Glu226 is substituted by a conserved Asp, and finally, most homologs show an Asn residue instead of Thr287 (Fig. 6). Therefore, as the retraction of the cap domain must be performed to allow substrate entrance, residues Phe91, Tyr190 and Tyr199 from the catalytic domain, which are located close to the catalytic triad, seem essential for substrate specificity (Fig. 5K).

## DISCUSSION

In this study, the microbial community of the *Sorghum bicolor* rhizosphere was exposed to a chemical treatment prior to eDNA extraction to construct a metagenomic library. Plant roots can secrete exudates composed of a large variety of compounds into the soil, some of which may play important roles in the rhizosphere (32, 33), and with effects that involve multiple targets, including soil microorganisms. This is why an amendment of the soil with technical cashew nut shell liquid (tCNSL), containing a mixture of phenolic compounds with long aliphatic side chains (up to C22:0), was carried out directly in the rhizosphere and, later, for three weeks under control laboratory conditions, as we were interested in screening lipolytic-like activity. By applying metagenomics techniques we retrieved an esterase, EH_0_, highly similar (99% identity) to the predicted α/β hydrolase from the genome of *S. pruni* (acc. nr. WP_066587239). The most homologous, functionally characterized protein is actually P95125.1, a carboxylic ester hydrolase LipN from *Mycobacterium tuberculosis* H37Rv, which shows only 41% amino acid sequence identity with EH_0_. That said, given the high identity of WP_066587239 and EH_0_ we expect both hydrolases having similar properties, yet to be experimentally confirmed. Indeed, we observed that there are only three changes in their sequences, which are located on the outside and in loops away from the key residues and the dimerisation interface (Fig. S2).

EH_0_ was classified within the previously described hormone-sensitive lipase (HDL) type IV family, that is one of the at least 35 families and 11 true lipase subfamilies known to date (10, 23, 34). This family is reported to contain ester hydrolases with relative SASA values ranging from 0 to 10% and high levels of substrate specificity (19). Note that SASA, computed as a (dimensionless) percentage (0−1 or 0-100) of the ligand SASA in solution (19), is a parameter that inspect the solvent exposure of the cavity containing the catalytic triad and the capacity of a cavity to retain/stabilize a substrate (19). For example, a SASA of 40% (over 100%) implies that 40% of the surface is accessible to the solvent, which facilitates the escape of the substrate to the bulk solvent; this is the case of enzymes with active site in the surface where the catalytic triad is highly exposed. By contrast, enzymes that has a larger but almost fully occluded site that can better maintain and stabilize the substrate inside the cavity, are characterized by relative SASA values of approximately 0−10%. This is the case of EH_0_, which has a SASA of 5.07% because a large but almost fully occluded active site, an architecture that is known to better maintain and stabilize a higher number of substrates inside the cavity (19). Indeed, this enzyme houses a very long and narrow catalytic pocket, where helix α7 is very close to the catalytic triad with residue Glu226, making a direct hydrogen bond to the catalytic nucleophile Ser161. Therefore, it appeared clear that the cap domain must retract to allow substrate entrance to the active site. The movement of this domain was modeled by combining X-ray diffraction data with molecular dynamics simulation through the ensemble refinement procedure. This strategy showed a broad range of putative conformations at the cap domain, with Pro46 and Pro47 likely acting as hinges conferring a high plasticity to the N-terminal region of the cap. Remarkably, the presence of a number of prolines at this region particularly these two sequential prolines, is a unique feature of EH_0_ compared to its homologs, and other substrate-promiscuous members of the family IV (19). Mutational analysis confirmed the role of one of these prolines in the access and hydrolysis of large and voluminous substrates and thus in the increase in the substrate promiscuity level.

The above structural features differ from those of other substrate promiscuous family IV esterases, tested over the same set of ester substrates, namely, EH_1AB1_ (31), and EH_3_ (35, 36) capable of hydrolyzing a similar number of esters. The comparison of EH_1AB1_ and EH_3_ with EH_0_, shows major differences related to the *lid* (Fig. 7A). Whereas EH_1AB1_ and EH_3_ show large and wide catalytic pockets with two possible access to the binding site (Fig. 7B and C), EH_0_ has a unique, narrow and long entrance to the catalytic active site, as said before (Fig. 7D). Therefore, in the case of EH_0_ only a structural rearrangement of the cap domain would allow its adaptation to all different substrates, which likely implies that the cap domain of EH_0_ exhibits more flexibility than those from EH_1AB1_ and EH_3_. This is consistent with the fact that of the 80 esters that the three enzymes together are able to hydrolyze, 61 (or 76%) were common to all three, and that EH_0_ was the only one able to hydrolyze such bulky substrates as 2,4-dichlorobenzyl 2,4-dichlorobenzoate or diethyl-2,6-dimethyl 4-phenyl-1,4-dihydro pyridine-3,5-dicarboxylate.

**FIG 7.**
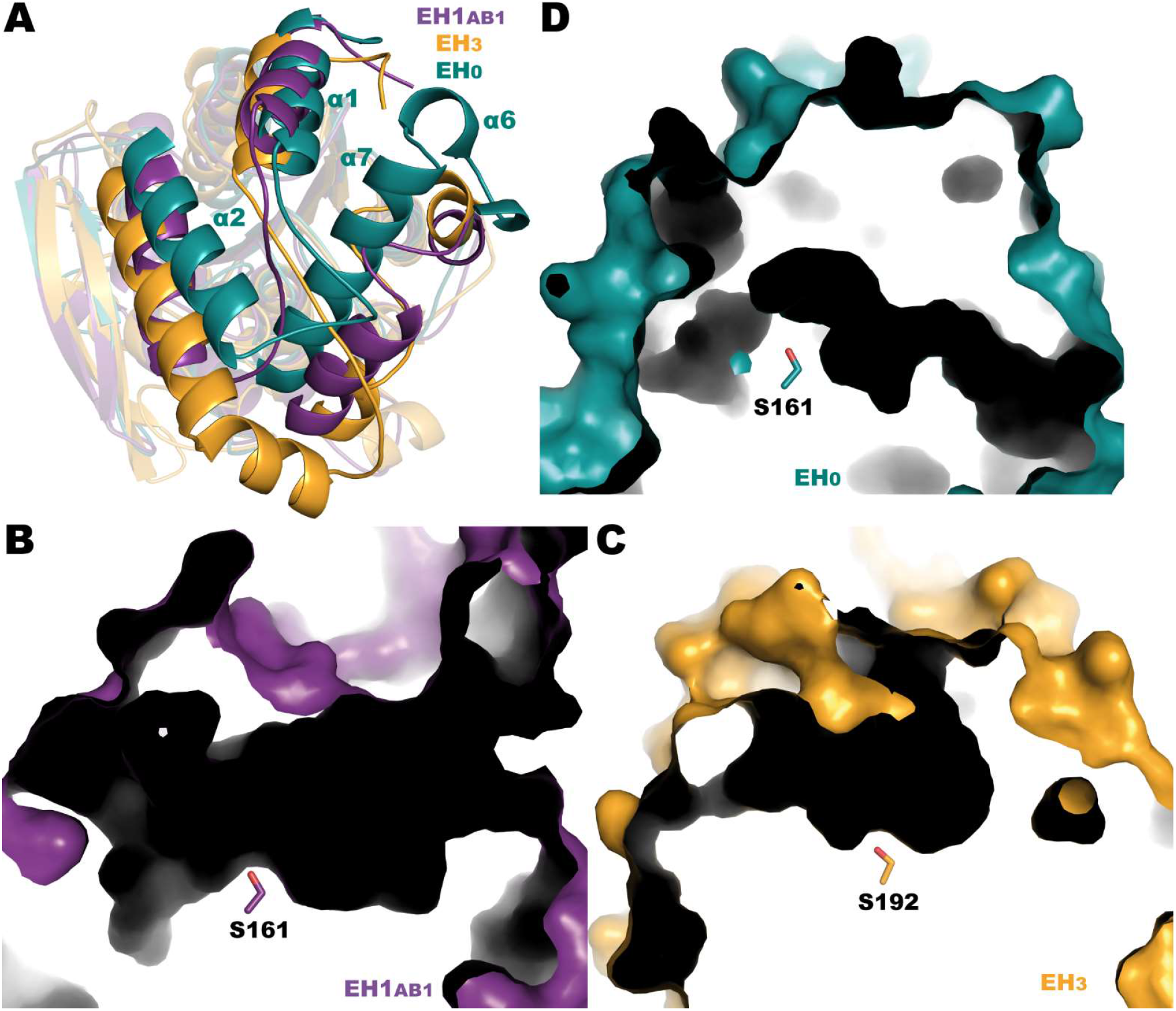
A) Superposition of EH_0_ (teal), EH_1AB1_ (purple) and EH_3_ (orange). Most differences are found in the cap domain. Catalytic cavities of EH_1AB1_ (B), EH_3_ (C) and EH_0_ (D).

Biochemical characterization of the novel esterase also revealed that the activity or EH_0_ was in most cases stimulated in the presence of 10% organic solvents, particularly in 10% methanol and DMSO. Such activation is also a characteristic of some lipases. For example, the analysis of the lipase from *Thermus thermophilus* revealed that although the overall structure was maintained stable with or without polar organic solvent, the lid region was more flexible in the presence of the later. The flexible lid facilitates the substrate to access the catalytic site inside the lipase and the lipase displays enhanced activity in the presence of a polar organic solvent (37). The use of organic solvents offers more advantages over canonical aqueous biocatalysis for various reasons: higher solubility of hydrophobic substrates, minor risk of contamination, and higher thermal stability (38–40). EH_0_ has a potential advantage in applications that require alkaline conditions due to its ability to act at the optimal pH of 9.0. Temperature-controlled tests indicated a mesophilic/slightly thermophilic profile of the esterase as expected from the original habitat at moderate temperatures. In addition, the structure of EH_0_ is similar (31-33% identity) to those of extremophiles, namely, *Pyrobaculum* (3ZWQ) and *Sulfolobus* (5LK6) species.

In summary, the present study evaluated the conformational plasticity of the cap domain in members of Family IV, and the role of several non-conserved prolines as putative structural factor regulating their broader substrate specificity compared to other members of the 35 families and 11 true lipase subfamilies reported (10). This high molecular flexibility is markedly different to that found in other family IV esterases and a family VIII β-lactamase fold hydrolase (EH_7_) which has been recently shown to be highly substrate-promiscuous. In this case, the broad substrate specificity is given by the presence of a more opened and exposed S1 site having no steric hindrance for the entrance of substrates to the active site, and more flexible R1, R2 and R3 regions allowing the binding of a wide spectrum of substrates into the active site.

## CONCLUSIONS

Activity-based metagenomics approach was used to study the microbial enzyme diversity in rhizosphere soil of *Sorghum* plants amended with CNSL soil. A novel esterase was found, which possessed a broad substrate promiscuity in combination with a significant pH and solvent tolerance. This work is crucial for deciphering structural markers responsible for the outstanding broad substrate specificity of EH_0_. Indeed, this work further provides important insights into the role of cap domains and its contribution to the diverse selectivity profiles and thus versatility of family IV esterases/lipases toward the conversion of multiple substrates.

## MATERIALS AND METHODS

### Plant material and outdoor seed germination

Seeds of *Sorghum bicolor* genotype BTx623 were obtained from the Agricultural Research Service of United States Department of Agriculture (USDA) (Georgia, US) as a gift. Field soil was sampled from the Henfaes Research Centre (53°14’21.0"N, 4°01’06.5"W, Gwynedd, Wales) in September 2014. The soil sample was composed of a mixture of five topsoil samples collected from randomly selected positions in the field. The soil was air-dried, mixed thoroughly, and stored at room temperature for use in subsequent experiments. Two L pots were filled with soil, and two seeds of *S. bicolor* BTx623 were planted per pot. Plants were cultivated in a greenhouse at 20 °C, and the soil moisture content was maintained with tap water.

### Enrichment with CNSL

Three grams of technical cashew nut shell liquid (tCNSL) dissolved in 70% ethanol was added to a pot of 20-day-old plants and thoroughly mixed with the soil. After 60 days, plants were pulled out of the pot, and the soil was shaken off; samples of rhizosphere soil attached to the plant roots were then brushed off and collected. tCNSL was provided by the BioComposites Centre at Bangor University (Wales, UK). Three biological replicates of laboratory microcosm enrichment were set up in conical 1 L Erlenmeyer flasks by mixing 10 g of the collected rhizosphere soil with 300 mL of sterile Murashige Skoog Basal medium (Sigma) and 10 mg/L cycloheximide. tCNSL was dissolved in 70% ethanol and added to the medium to a final concentration of 0.1 g/L; flasks were incubated at 20 °C in an orbital shaker. 50 g of oil slurry was sampled every seven days, and fresh tCNSL-containing (0.1 g/kg) medium was added to replace the volume of the medium.

### Extraction of DNA and generation of metagenomic library

Samples collected after 3 weeks of flask microcosm enrichment were used for the construction of fosmid metagenomic libraries. Environmental DNA was extracted using the Meta-G-Nome DNA Isolation Kit (Epicenter Biotechnologies; WI, USA) according to the manufacturer’s instructions. Briefly, 50 ml of the soil suspension from the flask enrichment was centrifuged at 400 × g for 5 min. The supernatant was filtered through 0.45 μm and 0.22 μm membrane filters. This procedure was repeated with the initial soil sample four times: the remaining soil was resuspended in PBS and centrifuged, and the supernatant was filtered as before. Filters were combined, and the sediment on the filter was resuspended in extraction buffer and collected. DNA extraction was carried out according to the protocol described by the manufacturer. The quality of the extracted DNA was evaluated on agarose gel and quantified with the Quant-iT dsDNA Assay Kit (Invitrogen) on a Cary Eclipse fluorimeter (Varian/Agilent) according to the manufacturer’s instructions. The extracted metagenomic DNA was used to prepare two different metagenomic fosmid libraries using the CopyControl™ Fosmid Library Production Kit (Epicenter). DNA was end-repaired to generate blunt-ended, 5’-phosphorylated double-stranded DNA using reagents included in the kit according to the manufacturer’s instructions. Subsequently, fragments of 30-40 kb were selected by electrophoresis and recovered from a low melting point agarose gel using GELase 50X buffer, and GELase enzyme preparation was also included in the kit. Nucleic acid fragments were then ligated to the linearized CopyControl pCC2FOS vector in a ligation reaction performed at room temperature for 4 hours, according to the manufacturer’s instructions. After in vitro packaging into phage lambda (MaxPlax™ Lambda Packaging Extract, Epicenter), the transfected phage T1-resistant EPI300™-T1R *E. coli* cells were spread on Luria-Bertani (LB) agar medium (hereinafter, unless mentioned otherwise, the agar content was 1.5% w/vol) containing 12.5 μg/ml chloramphenicol and incubated at 37 °C overnight to determine the titer of the phage particles. The resulting library, SorRhizCNSL3 W, has an estimated titer of 1.5×10^6^ clones. For long-term storage, the library was plated onto solid LB medium with 12.5 μg/ml chloramphenicol, and after overnight growth, colonies were washed off from the agar surface using LB broth with 20% (v/v) sterile glycerol, and aliquots were stored at −80 °C.

### Screening metagenomic libraries: agar-based methods

Fosmid clones obtained by plating the constructed libraries on LB agar plates were arrayed in 384-microtiter plates (1 clone/well) or alternatively in 96-microtiter plates (pools of approximately 40 clones/well) containing LB medium and chloramphenicol (12.5 μg/ml). The plates were incubated at 37 °C overnight, and the day after replication, the plates were produced and used in the screening assay. Glycerol (20% v/v, final concentration) was added to the original plates, which were stored at −80 °C. Gel diffusion and colorimetric assays were adopted for the screening of the desired activities. The detection of lipase/esterase activity was carried out on LB agar supplemented with chloramphenicol (12.5 μg/ml), fosmid autoinduction solution (2 ml/L) (Epicenter), and 0.3% (v/v) tributyrin emulsified with gum arabic (2:1, v/v) by sonication. The previously prepared microtiter plates were printed on the surface of large (22.5 cm × 22.5 cm) LB agar plates using 384-pin polypropylene replicators and incubated for 18-48 hours at 37 °C. Lipolytic activity was identified as a clear zone around the colonies where tributyrin was hydrolyzed (12).

### Extraction of fosmids, DNA sequencing and annotation

The fosmid DNA of the positive clone was extracted using the QIAGEN Plasmid purification kit (QIAGEN). To reduce the host chromosomal *E. coli* DNA contamination, the sample was treated with ATP-dependent exonuclease (Epicenter). The purity and approximate size of the cloned fragment were assessed by agarose gel electrophoresis after endonuclease digestion simultaneously with *BamHI* and *XbaI* (New England Biolabs, in NEBuffer 3.1 at 37° for 1 hour using 1 U of enzyme per 1 μg DNA). DNA concentration was quantified using the Quant-iT dsDNA Assay Kit (Invitrogen), and DNA sequencing was then outsourced to Fidelity Systems (NJ, USA) for shotgun sequencing using the Illumina MiSeq platform. GeneMark software (41) was employed to predict protein coding regions from the sequences of each assembled contig, and deduced amino acid sequences were annotated via BLASTP and the PSI-BLAST tool (42).

### Cloning, expression and purification of proteins

The selected nucleotide sequence was amplified by PCR using Herculase II Fusion Enzyme (Agilent, USA) with specific oligonucleotide primer pairs incorporating p15TV-L adapters. The corresponding fosmid was used as a template to amplify the target genes. The primers used to amplify the esterase gene characterized in this study were as follows: EH_0F_, TTGTATTTCCAGGGCATGACCGAGCTCTTCGTCCGC; EH_0R_, CAAGCTTCGTCATCATGCCGCCGCCTGTGCCATC. PCR products were visualized on a 1% TAE agarose gel and purified using the Nucleospin PCR Clean-up Kit (Macherey-Nagel) following the manufacturer’s instructions. Purified PCR products were cloned into the p15TV-L vector, transformed into *E. coli* NovaBlue GigaSingles™ Competent Cells (Novagen, Germany) and plated on LB agar with 100 μg/ml ampicillin. The correctness of the DNA sequence was then verified by Sanger sequencing at Macrogen Ltd. (Amsterdam, The Netherlands). 3-D models of the proteins were generated by Phyre2. The intensive mode attempts to create a complete full-length model of a sequence through a combination of multiple template modeling and simplified ab initio folding simulation (43). The nucleotide and amino acid sequences of the selected nucleotide sequences are available GenBank under ref. MK791218. For recombinant protein expression, the plasmids were transformed into *E. coli* BL21(DE3) cells and subsequently plated on LB agar with 100 μg/ml ampicillin. To confirm the esterase activity of recombinant proteins, *E. coli* clones harboring the recombinant plasmid were streaked onto LB agar plates containing 0.5% tributyrin and 0.5 mM IPTG, or purified enzymes were spotted directly on the agar. The plates were then incubated at 37 °C overnight and visually inspected for the presence of signs of substrate degradation. *E. coli* clones were grown at 37 °C to an absorbance of 0.8 at 600 nm, induced with 0.5 mM isopropyl-β-D-galactopyranoside (IPTG), and allowed to grow overnight at 20 °C with shaking. Cells were harvested by centrifugation at 5000 × g for 30 min at 4 °C. For purification of recombinant protein, the following protocol was applied. Cell pellets were resuspended in cold binding buffer (50 mM HEPES pH 7.5, 400 mM NaCl, 5% glycerol, 0.5% Triton-X, 6 mM imidazole pH 7.5, 1 mM β-mercaptoethanol, 0.5 mM PMSF) and extracted by sonication. The lysates were then centrifuged at 22,000 × g for 30 min at 4 °C, and the supernatant was purified by affinity chromatography using Ni-NTA His-bind resin (Novagen). The column packed with the resin was equilibrated with binding buffer, and after the addition of supernatant, it was washed with 6 volumes of wash buffer (50 mM HEPES pH 7.5, 400 mM NaCl, 5% glycerol, 0.5% Triton-X, 26 mM imidazole pH 7.5, 1 mM β-mercaptoethanol, 0.5 mM PMSF) to remove nonspecifically bound proteins. His-tagged proteins were then eluted with elution buffer (50 mM HEPES pH 7.5, 400 mM NaCl, 5% glycerol, 0.5% Triton-X, 266 mM imidazole pH 7.5, 1 mM β-mercaptoethanol, 0.5 mM PMSF). The size and purity of the proteins were estimated by SDS–PAGE. Protein solutions were desalted through the Amicon Ultra15 10K centrifugal filter device. Protein concentrations were determined using the Bradford Reagent (Sigma) and the BioMate™ 3S spectrophotometer (Thermo Scientific, USA).

### Biochemical assays

Hydrolytic activity was determined by measuring the amount of *p*-nitrophenol released by catalytic hydrolysis of *p*-nitrophenyl (*p*-NP) esters through a modified method of Gupta et al. (44). Stock solutions of *p*-NP esters (100 mM *p*-NP acetate, 100 mM *p*-NP butyrate, 20 mM *p*-NP dodecanoate, and 20 mM *p*-NP palmitate) were prepared in DMSO/acetonitrile (1:1 v/v). Unless stated otherwise, the enzymatic assay was performed under standard conditions in a 1 ml reaction (50 mM potassium phosphate buffer pH 7.0, 0.3% (v/v) Triton X-100, 1 mM substrate) under agitation in a water bath at 50 °C until complete substrate solubilization (note: solvents were present in the assays at a concentration of 1% for 100 mM-stocks of *p*-NP butyrate and acetate, or 5% for 20 mM-stocks of *p*-NP dodecanoate and palmitate). After that, the multititre plate was pre-incubated at 30 °C for 5 minutes in a BioMate™ 3S spectrophotometer (Thermo Scientific, USA), which was set up at this temperature, and then an appropriate volume of purified enzyme containing 0.25 μg was added to start the reaction. The reaction mixture was incubated at 30 °C for 5 min and then measured at 410 nm in BioMate™ 3S spectrophotometer (Thermo Scientific, USA). The incubation time for *p*-NP dodecanoate and *p*-NP palmitate was extended to 15 minutes. All experiments were performed in triplicate, and a blank with denatured enzyme was included. The concentration of product was calculated by linear regression equation given on the standard curve performed by the reference compound *p*-nitrophenol (Sigma). One unit of enzyme activity was defined as 1 μmol of *p*-nitrophenol produced per minute under the assay conditions.

Kinetic parameters were determined under standard conditions and calculated by nonlinear regression analysis of raw data fit to the Michaelis–Menten function using GraphPad Prism software (version 6.0). For the kinetics with *p*-NP butyrate and acetate the concentrations were set up to 0.1, 0.5, 1, 2 and 10 mM, using stock concentrations of 100 mM in DMSO/acetonitrile (a maximum concentration of 5% DMSO/acetonitrile, was used in the assay). For *p*-NP dodecanoate, a concentration of 0.05, 0.1, 0.5, 1, 2 and 3 mM was used, with stock concentration of 20 mM substrate (a maximum concentration of 7.5% DMSO/acetonitrile, was used in the assay). Raw data, and information about precision and calculations are provided in Table S6.

The optimal pH for enzyme activity was evaluated with *p*-NP butyrate by performing the assay in different buffers, specifically 20 mM sodium acetate buffer (pH 4.0), sodium citrate buffer (pH 5.5), potassium phosphate buffer (pH 7.0), and Tris-HCl (pH 8.0-9.0). The enzyme reactions were stopped by adding 1 ml of cold stop solution (100 mM K-phosphate buffer pH 7.0, 10 mM EDTA) to neutralize the pH and avoid changes in the equilibrium between *p*-nitrophenol and the deprotonated form *p*-nitrophenoxide, which would result in a decrease in absorption at the applied wavelength of 410 nm (45). The optimal enzymatic temperature was investigated with *p*-NP butyrate by performing the hydrolytic assay at different temperatures under standard conditions (see above). To determine the denaturation temperature, circular dichroism (CD) spectra were acquired between 190 and 270 nm with a Jasco J-720 spectropolarimeter equipped with a Peltier temperature controller in a 0.1-mm cell at 25 °C. The spectra were analyzed, and melting temperature (T_m_) values were determined at 220 nm between 10 and 85 °C at a rate of 30 °C per hour in 40 mM HEPES buffer at pH 7.0. CD measurements were performed at pH 7.0 and not at the optimal pH (8.5-9.0) to ensure protein stability. A protein concentration of 0.5 mg∙mL^−1^ was used. T_m_ (and the standard deviation of the linear fit) was calculated by fitting the ellipticity (mdeg) at 220 nm at each of the different temperatures using a 5-parameter sigmoid fit with SigmaPlot 13.0.

Stability in organic solvents was assayed with *p*-NP butyrate under standard conditions in the presence of 10-20-40-60% (v/v) of the water-miscible organic solvents ethanol, methanol, isopropanol, acetonitrile, DMSO, and a mixture acetonitrile/DMSO (50% each). The effect of cations was investigated with *p*-NP butyrate under standard conditions by the addition of MgCl_2_, CuCl_2_, FeCl_3_, CoCl_2_, CaCl_2_, MnCl_2_, and ZnSO_4_ at concentrations in the range of 1 to 10 mM. In all cases, the measured values were then expressed as the relative activity in comparison to the control reaction performed under standard conditions.

The hydrolysis of esters other than *p*-NP esters, including bis(2-hydroxyethyl)-terephthalate (BHET), was assayed using a pH indicator assay in 384-well plates at 30 °C and pH 8.0 in a Synergy HT Multi-Mode Microplate Reader in continuous mode at 550 nm over 24 h (extinction coefficient of phenol red, 8450 M^−1^cm^−1^). The acid produced after ester bond cleavage by the hydrolytic enzyme induced a color change in the pH indicator that was measured at 550 nm. The experimental conditions were as detailed previously (35), with the absence of activity defined as at least a twofold background signal. Briefly, the reaction conditions for 384-well plates (ref. 781162, Greiner Bio-One GmbH, Kremsmünster, Austria) were as follows: protein, 0.2-2.0 μg per well; ester, 20 mM; T: 30 °C; pH, 8.0 (5 mM 4-(2-hydroxyethyl)-1-piperazinepropanesulfonic acid (EPPS) buffer, plus 0.45 Phenol Red^®^); reaction volume, 40 μl. The reactions were performed in triplicate, and datasets were collected from a Synergy HT Multi-Mode Microplate Reader with Gen5 2.00 software (Biotek). One unit (U) of enzyme activity was defined as the amount of enzyme required to transform 1 μmol of substrate in 1 min under the assay conditions. Raw data, and information about precision and calculations are provided in Table S7.

### Crystallization and X-ray structure determination of EH_0_

Initial crystallization conditions were explored by high-throughput techniques with a NanoDrop robot (Innovadyne Technologies) using 24 mg∙mL^−1^ protein concentrations in HEPES (40 mM, pH 7, NaCl 50 mM), protein reservoir ratios of 1:1, 1.5:1 and 2:1, and commercial screens Crystal Screen I and II, SaltRx, Index (Hampton Research), JBScreen Classic, JBScreen JCSG, JBScreen PACT (Jena Bioscience). Further optimizations were carried out, and bar-shaped crystals of EH_0_ were grown after one day of mixing 1.2 μL of a mixture of protein (1 μL, 24 mg∙mL^−1^) and seeds (0.2 μL, 1:100) with guanidine hydrochloride (0.2 μL, 0.1 M) and reservoir (0.5 μL, 11% polyethylene glycol 8000, 100 mM Bis-Tris pH 5.5, 100 mM ammonium acetate). For data collection, crystals were transferred to cryoprotectant solution consisting of mother liquour and glycerol (20% (v/v)) before being cooled in liquid nitrogen. Diffraction data were collected using synchrotron radiation on the XALOC beamline at ALBA (Cerdanyola del Vallés, Spain). Diffraction images were processed with XDS (46) and merged using AIMLESS from the CCP4 package (47). The crystal was indexed in the P2_1_2_1_2_1_ space group, with two molecules in the asymmetric unit and 62% solvent content within the unit cell. The data collection statistics are given in Table S4. The structure of EH_0_ was solved by molecular replacement with MOLREP (48) using the coordinates from Est8 as a template (PDB Code 4YPV). Crystallographic refinement was performed using the program REFMAC (49) within the CCP4 suite, with NCS (noncrystallography symmetry) medium restraints and excluding residues 17-36. The free *R*-factor was calculated using a subset of 5% randomly selected structure-factor amplitudes that were excluded from automated refinement. Subsequently, heteroatoms were manually built into the electron density maps with Coot8 (50), and water molecules were included in the model, which, combined with more rounds of restrained refinement, reached the *R* factors listed in Table S4. The figures were generated with PyMOL. The crystallographic statistics of EH_0_ are listed in Table S4. To extract dynamical details from the X-ray data, the coordinates of EH_0_ were first refined using PHENIX (51) and then were used as input models for a time-averaged molecular dynamics refinement as implemented in the Phenix.ensemble-refinement routine, which was performed as described previously (52).

### Codes and accession numbers

The sequence encoding EH_0_ was deposited in GenBank with the accession number MK791218. The atomic coordinates and structure factors for the EH_0_ structure have been deposited in the RCSB Protein Data Bank with accession codes 7ZR3.

## SUPPLEMENTARY MATERIAL

Supplemental material is available online only.

## ACKNOWLEDGMENTS

This study was conducted under the auspices of the FuturEnzyme Project funded by the European Union’s Horizon 2020 Research and Innovation Programme under Grant Agreement No. 101000327. We also acknowledge financial support under Grants PID2020-112758RB-I00 (M.F.), PDC2021-121534-I00 (M.F.), and PID2019-105838RB-C33 (J.S-A.) from the Ministerio de Ciencia e Innovación, Agencia Estatal de Investigación (AEI) (Digital Object Identifier 10.13039/501100011033), Fondo Europeo de Desarrollo Regional (FEDER) and the European Union (“NextGenerationEU/PRTR”), and Grant 2020AEP061 (M.F.) from the Agencia Estatal CSIC. C. Coscolín thanks the Ministerio de Economía y Competitividad and FEDER for a PhD fellowship (Grant BES-2015-073829). P.N.G. acknowledges the Sêr Cymru programme partly funded by ERDF through the Welsh Government for the support of the project BioPOL4Life, the project ‘Plastic Vectors’ funded by the Natural Environment Research Council UK (NERC), Grant No. NE/S004548/N and the Centre for Environmental Biotechnology Project co-funded by the European Regional Development Fund (ERDF) through the Welsh Government. The authors acknowledge David Almendral and Ruth Matesanz for supporting the circular dichroism analysis, and Jose L. Gonzalez-Alfonso and Francisco J. Plou for supporting the HPLC analysis. We thank the staff of the Synchrotron Radiation Source at Alba (Barcelona, Spain) for assistance at the BL13-XALOC beamline.

We declare no conflict of interest.

P.N.G., M.F. and J.S-A. developed the conceptual framework, M.D., C.C., T.N.C. and H.T. developed the experimental design, I.C-R. and J.S-A. performed the crystallization and X-ray structure determinations, analysis and interpretation, M.D., M.F. and P.N.G. interpreted the experimental data, P.N.G., M.F. and J.S-A. wrote the initial manuscript. All authors edited the manuscript. All authors read and approved the final manuscript.

## References

1. Intasian P, Prakinee K, Phintha A, Trisrivirat D, Weeranoppanant N, Wongnate T, Chaiyen P. 2021. Enzymes, in vivo biocatalysis, and metabolic engineering for enabling a circular economy and sustainability. Chem Rev 121:10367–10451.

2. Ferrer M, Méndez-García C, Bargiela R, Chow J, Alonso S, García-Moyano A, Bjerga GEK, Steen IH, Schwabe T, Blom C, Vester J, Weckbecker A, Shahgaldian P, De Carvalho CCCR, Meskys R, Zanaroli G, Glöckner FO, Fernández-Guerra A, Thambisetty S, De La Calle F, Golyshina O V., Yakimov MM, Jaeger KE, Yakunin AF, Streit WR, McMeel O, Calewaert JB, Tonné N, Golyshin PN. 2019. Decoding the ocean’s microbiological secrets for marine enzyme biodiscovery. FEMS Microbiol Lett 366:1–7.

3. Schloss PD, Handelsman J. 2006. Toward a census of bacteria in soil. PLoS Comput Biol 2:0786–0793.

4. Handelsman J. 2005. Metagenomics: Application of genomics to uncultured microorganisms. Microbiol Mol Biol Rev 69:195–195.

5. Barzkar N, Sohail M, Tamadoni Jahromi S, Gozari M, Poormozaffar S, Nahavandi R, Hafezieh M. 2021. Marine bacterial esterases: Emerging biocatalysts for industrial applications. Appl Biochem Biotechnol 193:1187–1214.

6. Martínez-Martínez M, Alcaide M, Tchigvintsev A, Reva O, Polaina J, Bargiela R, Guazzaroni ME, Chicote Á, Canet A, Valero F, Eguizabal ER, Guerrero M del C, Yakunin AF, Ferrer M. 2013. Biochemical diversity of carboxyl esterases and lipases from lake arreo (Spain): A metagenomic approach. Appl Environ Microbiol 79:3553–3562.

7. Jaeger KE, Eggert T. 2002. Lipases for biotechnology. Curr Opin Biotechnol 13:390–397.

8. Jochens H, Hesseler M, Stiba K, Padhi SK, Kazlauskas RJ, Bornscheuer UT. 2011. Protein engineering of α/β-hydrolase fold enzymes. ChemBioChem 12:1508–1517.

9. De Godoy Daiha K, Angeli R, De Oliveira SD, Almeida RV. 2015. Are lipases still important biocatalysts? A study of scientific publications and patents for technological forecasting. PLoS One 10:1–20.

10. Bauer TL, Buchholz PCF, Pleiss J. 2020. The modular structure of α/β-hydrolases. FEBS J 287:1035–1053.

11. Ferrer M, Martínez-Martínez M, Bargiela R, Streit WR, Golyshina O V., Golyshin PN. 2016. Estimating the success of enzyme bioprospecting through metagenomics: Current status and future trends. Microb Biotechnol 9:22–34.

12. Placido A, Hai T, Ferrer M, Chernikova TN, Distaso M, Armstrong D, Yakunin AF, Toshchakov S V., Yakimov MM, Kublanov I V., Golyshina O V., Pesole G, Ceci LR, Golyshin PN. 2015. Diversity of hydrolases from hydrothermal vent sediments of the Levante Bay, Vulcano Island (Aeolian archipelago) identified by activity-based metagenomics and biochemical characterization of new esterases and an arabinopyranosidase. Appl Microbiol Biotechnol 99:10031–10046.

13. Tchigvintsev A, Tran H, Popovic A, Kovacic F, Brown G, Flick R, Hajighasemi M, Egorova O, Somody JC, Tchigvintsev D, Khusnutdinova A, Chernikova TN, Golyshina O V., Yakimov MM, Savchenko A, Golyshin PN, Jaeger KE, Yakunin AF. 2015. The environment shapes microbial enzymes: five cold-active and salt-resistant carboxylesterases from marine metagenomes. Appl Microbiol Biotechnol 99:2165–2178.

14. Lopez-Lopez O, Cerdan ME, Gonzalez Siso MI. 2014. New Extremophilic Lipases and Esterases from Metagenomics. Curr Protein Pept Sci 15:445–455.

15. Lee MH, Hong KS, Malhotra S, Park JH, Hwang EC, Choi HK, Kim YS, Tao W, Lee SW. 2010. A new esterase EstD2 isolated from plant rhizosphere soil metagenome. Appl Microbiol Biotechnol 88:1125–1134.

16. Faoro H, Glogauer A, Souza EM, Rigo LU, Cruz LM, Monteiro RA, Pedrosa FO. 2011. Identification of a new lipase family in the Brazilian Atlantic Forest soil metagenome. Environ Microbiol Rep 3:750–755.

17. Hult K, Berglund P. 2007. Enzyme promiscuity: mechanism and applications. Trends Biotechnol 25:231–238.

18. Alcaide M, Tornés J, Stogios PJ, Xu X, Gertler C, Leo RDI, Bargiela R, Lafraya Á, Guazzaroni ME, López-Cortés N, Chernikova TN, Golyshina O V., Nechitaylo TY, Plumeier I, Pieper DH, Yakimov MM, Savchenko A, Golyshin PN, Ferrer M. 2013. Single residues dictate the co-evolution of dual esterases: MCP hydrolases from the α/β hydrolase family. Biochem J 454:157–166.

19. Martínez-Martínez M, Coscolín C, Santiago G, Chow J, Stogios PJ, Bargiela R, Gertler C, Navarro-Fernández J, Bollinger A, Thies S, Méndez-García C, Popovic A, Brown G, Chernikova TN, García-Moyano A, Bjerga GEK, Pérez-García P, Hai T, Del Pozo M V., Stokke R, Steen IH, Cui H, Xu X, Nocek BP, Alcaide M, Distaso M, Mesa V, Peláez AI, Sánchez J, Buchholz PCF, Pleiss J, Fernández-Guerra A, Glöckner FO, Golyshina O V., Yakimov MM, Savchenko A, Jaeger KE, Yakunin AF, Streit WR, Golyshin PN, Guallar V, Ferrer M. 2018. Determinants and prediction of esterase substrate promiscuity patterns. ACS Chem Biol 13:225–234.

20. Höppner A, Bollinger A, Kobus S, Thies S, Coscolín C, Ferrer M, Jaeger KE, Smits SHJ. 2021. Crystal structures of a novel family IV esterase in free and substrate-bound form. FEBS J 288:3570–3584.

21. Kumar PP, Paramashivappa R, Vithayathil PJ, Rao PVS, Rao AS. 2002. Process for isolation of cardanol from technical cashew (*Anacardium occidentale* l.) Nut shell liquid. J Agric Food Chem 50:4705–4708.

22. Reyes-Duarte D, Ferrer M, García-Arellano H. 2012. Functional-based screening methods for lipases, esterases, and phospholipases in metagenomic libraries. Lipases and Phospholipases 861:101–113.

23. Arpigny JL, Jaeger KE. 1999. Bacterial lipolytic enzymes: Classification and properties. Biochem J 343:177–183.

24. Yoshida S, Hiraga K, Takehana T, Taniguchi I, Yamaji H, Maeda Y, Toyohara K, Miyamoto K, Kimura Y, Oda K. 2016. A bacterium that degrades and assimilates poly(ethylene terephthalate). Sci (American Assoc Adv Sci 351:1196–1199.

25. Bollinger A, Thies S, Knieps-Grünhagen E, Gertzen C, Kobus S, Höppner A, Ferrer M, Gohlke H, Smits SHJ, Jaeger KE. 2020. A novel polyester hydrolase from the marine bacterium *Pseudomonas aestusnigri* – Structural and functional insights. Front Microbiol 11:1–16.

26. Pereira MR, Maester TC, Mercaldi GF, de Macedo Lemos EG, Hyvönen M, Balan A. 2017. From a metagenomic source to a high-resolution structure of a novel alkaline esterase. Appl Microbiol Biotechnol 101:4935–4949.

27. Holm L. 2020. Using Dali for Protein Structure Comparison. Struct Bioinforma 2112:29–42.

28. Ding Y, Nie L, Yang XC, Li Y, Huo YY, Li Z, Gao Y, Cui HL, Li J, Xu XW. 2022. Mechanism and structural insights into a novel esterase, E53, isolated From *Erythrobacter longus*. Front Microbiol 12:1–12.

29. Palm GJ, Fernández-Álvaro E, Bogdanović X, Bartsch S, Sczodrok J, Singh RK, Böttcher D, Atomi H, Bornscheuer UT, Hinrichs W. 2011. The crystal structure of an esterase from the hyperthermophilic microorganism *Pyrobaculum calidifontis* VA1 explains its enantioselectivity. Appl Microbiol Biotechnol 91:1061–1072.

30. De Simone G, Menchise V, Alterio V, Mandrich L, Rossi M, Manco G, Pedone C. 2004. The crystal structure of an EST2 mutant unveils structural insights on the H group of the carboxylesterase/lipase family. J Mol Biol 343:137–146.

31. Alonso S, Santiago G, Cea-Rama I, Fernandez-Lopez L, Coscolín C, Modregger J, Ressmann AK, Martínez-Martínez M, Marrero H, Bargiela R, Pita M, Gonzalez-Alfonso JL, Briand ML, Rojo D, Barbas C, Plou FJ, Golyshin PN, Shahgaldian P, Sanz-Aparicio J, Guallar V, Ferrer M. 2020. Genetically engineered proteins with two active sites for enhanced biocatalysis and synergistic chemo- and biocatalysis. Nat Catal 3:319–328.

32. Bais HP, Weir TL, Perry LG, Gilroy S, Vivanco JM. 2006. The role of root exudates in rhizosphere interactions with plants and other organisms. Annu Rev Plant Biol 57:233–266.

33. Bertin C, Yang X, Weston LA. 2003. The role of root exudates and allelochemicals in the rhizosphere. Plant Soil 256:67–83.

34. Hitch TCA, Clavel T. 2019. A proposed update for the classification and description of bacterial lipolytic enzymes. PeerJ 2019.

35. Giunta CI, Cea-Rama I, Alonso S, Briand ML, Bargiela R, Coscolín C, Corvini PFX, Ferrer M, Sanz-Aparicio J, Shahgaldian P. 2020. Tuning the properties of natural promiscuous enzymes by engineering their nano-environment. ACS Nano 14:17652–17664.

36. Cea-Rama I, Coscolín C, Katsonis P, Bargiela R, Golyshin PN, Lichtarge O, Ferrer M, Sanz-Aparicio J. 2021. Structure and evolutionary trace-assisted screening of a residue swapping the substrate ambiguity and chiral specificity in an esterase. Comput Struct Biotechnol J 19:2307–2317.

37. Cao H, Nie K, Xu H, Xiong X, Krastev R, Wang F, Tan T, Liu L. 2016. Insight into the mechanism behind the activation phenomenon of lipase from *Thermus thermophilus* HB8 in polar organic solvents. J Mol Catal B Enzym 133:S400–S409.

38. Sellek GA, Chaudhuri JB. 1999. Biocatalysis in organic media using enzymes from extremophiles. Enzyme Microb Technol 25:471–482.

39. Zaks A, Klibanov AM. 1988. Enzymatic catalysis in nonaqueous solvents. J Biol Chem 263:3194–3201.

40. Sharma S, Kanwar SS. 2014. Organic solvent tolerant lipases and applications. Sci World J 2014.

41. Lukashin A V., Borodovsky M. 1998. GeneMark.hmm: New solutions for gene finding. Nucleic Acids Res 26:1107–1115.

42. Altschul SF, Madden TL, Schäffer AA, Zhang J, Zhang Z, Miller W, Lipman DJ. 1997. Gapped BLAST and PSI-BLAST: A new generation of protein database search programs. Nucleic Acids Res 25:3389–3402.

43. Kelley LA, Mezulis S, Yates CM, Wass MN, Sternberg MJE. 2015. The Phyre2 web portal for protein modeling, prediction and analysis. Nat Protoc 10:845–858.

44. Gupta N, Rathi P, Gupta R. 2002. Simplified para-nitrophenyl palmitate assay for lipases and esterases. Anal Biochem 311:98–99.

45. Bowers GN, McComb RB, Christensen RG, Schaffer R. 1980. High-purity 4-nitrophenol: purification, characterization, and specifications for use as a spectrophotometric reference material. Clin Chem 26:724–729.

46. Kabsch W. 2010. research papers XDS research papers. Acta Crystallogr Sect D Biol Crystallogr 66:125–132.

47. Evans PR, Murshudov GN. 2013. How good are my data and what is the resolution? Acta Crystallogr Sect D Biol Crystallogr 69:1204–1214.

48. Vagin A, Teplyakov A. 1997. MOLREP: an Automated Program for Molecular Replacement. J Appl Crystallogr 30:1022–1025.

49. Murshudov GN, Vagin AA, Dodson EJ. 1997. Refinement of Macromolecular Structures by the Maximum-Likelihood Method. Acta Crystallogr Sect D, Biol Crystallogr 53:240–255.

50. Emsley P, Lohkamp B, Scott WG, Cowtan K. 2010. Features and development of Coot. Acta Crystallogr Sect D Biol Crystallogr 66:486–501.

51. Liebschner D, Afonine P V., Baker ML, Bunkoczi G, Chen VB, Croll TI, Hintze B, Hung LW, Jain S, McCoy AJ, Moriarty NW, Oeffner RD, Poon BK, Prisant MG, Read RJ, Richardson JS, Richardson DC, Sammito MD, Sobolev O V., Stockwell DH, Terwilliger TC, Urzhumtsev AG, Videau LL, Williams CJ, Adams PD. 2019. Macromolecular structure determination using X-rays, neutrons and electrons: Recent developments in Phenix. Acta Crystallogr Sect D Struct Biol 75:861–877.

52. Tom Burnley B, Afonine P V., Adams PD, Gros P. 2012. Modelling dynamics in protein crystal structures by ensemble refinement. Elife 2012:1–29.

