## Supplementary Tables S1-7 and Figures S1-S2 for "The mobility of the cap domain is essential for the substrate promiscuity of a family IV esterase from sorghum rhizosphere microbiome"

Table S1. Specific activity of purified $EH_0$ against different substrates. The results are mean from three independent experiments, with SD within 5% in all cases.

| | Specific activity (U/g)[a] | | | |
|---|---|---|---|---|
| Substrate | $EH_0$ | $EH_{0E226A}$ | $EH_{0Y223A}$ | $EH_{0P46A}$ |
| 1-Napthyl acetate | 9,82E+03 | 2,48E+04 | 9,82E+03 | 2,88E+04 |
| 1-Naphthyl butyrate | 2,03E+03 | 3,33E+03 | 2,04E+03 | 1,50E+04 |
| Glyceryl triacetate | 8,40E+03 | 2,21E+04 | 8,40E+03 | 1,88E+04 |
| Glyceryl tripropionate | 2,94E+04 | 3,20E+04 | 2,93E+04 | 3,18E+04 |
| Glyceryl tributyrate | 6,61E+03 | 1,71E+04 | 6,65E+03 | 2,12E+04 |
| Glyceryl trioctanoate | 0 | 0 | 0 | 5,20E+00 |
| Methyl 2-bromobutyrate | 0 | 0 | 0 | 3,90E+00 |
| Hexyl acetate | 3,28E+03 | 1,88E+04 | 3,35E+03 | 2,92E+03 |
| Octyl acetate | 2,02E+02 | 1,81E+03 | 2,10E+02 | 1,94E+03 |
| Dodecanoyl acetate | 0 | 2,86E+01 | 8,50E+00 | 8,79E+01 |
| Pentadecyl acetate | 0 | 1,04E+01 | 9,10E+00 | 1,57E+02 |
| Ethyl acetate | 0 | 1,50E+01 | 6,90E+00 | 2,40E+02 |
| Ethyl propionate | 0 | 6,51E+01 | 3,45E+01 | 1,64E+03 |
| Ethyl butyrate | 9,80E+00 | 9,11E+01 | 9,70E+00 | 1,35E+02 |
| Ethyl hexanoate | 1,30E+04 | 2,37E+04 | 1,14E+04 | 2,68E+03 |
| Ethyl octanoate | 1,63E+03 | 5,92E+03 | 1,72E+03 | 2,65E+03 |
| Ethyl decanoate | 5,21E+01 | 1,11E+02 | 5,63E+01 | 1,74E+02 |
| Ethyl benzoate | 3,06E+01 | 9,18E+01 | 3,50E+01 | 2,10E+02 |
| (1*R*)-(-)-Menthyl acetate | 3,91E+01 | 1,28E+02 | 4,08E+01 | 1,11E+02 |
| (1*S*)-(+)-Menthyl acetate | 6,50E+00 | 7,94E+01 | 6,70E+00 | 6,31E+01 |
| Methyl (*R*)-(-)-mandelate | 7,80E+00 | 5,00E-01 | 8,00E+00 | 6,57E+01 |
| Methyl (*S*)-(+)-mandelate | 2,60E+01 | 1,04E+02 | 2,81E+01 | 7,88E+02 |
| Ethyl (*R*)-(+)-4-chloro-3-hydroxybutyrate | 5,20E+00 | 5,21E+01 | 5,50E+00 | 4,47E+02 |
| Ethyl (*S*)-(−)-4-chloro-3-hydroxybutyrate | 1,43E+02 | 3,19E+02 | 1,51E+02 | 2,75E+03 |
| (+)-Ethyl D-Lactate | 7,55E+01 | 4,56E+01 | 7,54E+01 | 5,69E+02 |
| (−)-Ethyl L-lactate | 1,30E+02 | 3,58E+02 | 1,36E+02 | 3,04E+03 |
| (+)-Methyl (*S*)-3-hydroxybutyrate | 5,99E+01 | 2,60E+01 | 6,18E+01 | 1,97E+02 |
| Benzoic acid, 4-formyl-, phenylmethyl ester | 2,93E+02 | 1,65E+03 | 2,99E+02 | 3,14E+02 |
| (-)-Methyl (*R*)-3-hydroxybutyrate | 1,11E+02 | 2,15E+02 | 1,15E+02 | 4,55E+02 |
| (1R)-(-)-Neomenthyl acetate | 3,32E+01 | 1,62E+02 | 3,15E+01 | 1,25E+02 |
| Propylparaben | 7,80E+00 | 3,58E+01 | 7,60E+00 | 2,67E+01 |
| Butylparaben | 4,60E+00 | 1,89E+01 | 4,60E+00 | 4,56E+01 |
| Methyl 3-hydroxybenzoate | 0 | 0 | 5,01E+01 | 2,54E+01 |
| Phthalic acid diethyl ester | 5,20E+00 | 2,02E+01 | 5,60E+00 | 5,86E+01 |
| (−)-Methyl (*R*)-3-hydroxyvalerate | 5,01E+01 | 1,95E+01 | 5,11E+01 | 2,40E+02 |
| (+)-Methyl (*S*)-3-hydroxyvalerate | 1,27E+02 | 9,76E+03 | 1,26E+02 | 2,51E+03 |
| Benzyl (*R*)-(+)-2-hydroxy-3-phenylpropionate | 2,30E+04 | 3,09E+04 | 2,30E+04 | 3,58E+04 |
| Benzylparaben | 1,12E+02 | 2,27E+02 | 1,15E+02 | 4,08E+02 |
| Methyl benzoate | 2,80E+01 | 6,50E+00 | 2,83E+01 | 1,61E+02 |
| Methyl butyrate | 3,30E+00 | 6,50E+00 | 3,20E+00 | 2,93E+01 |

| | | | | |
|---|---|---|---|---|
| Methyl decanoate | 7,81E+01 | 3,25E+02 | 8,02E+01 | 2,77E+02 |
| Methyl hexanoate | 3,18E+03 | 4,75E+03 | 3,19E+03 | 1,52E+04 |
| Methyl octanoate | 2,12E+03 | 3,87E+03 | 1,16E+03 | 7,26E+03 |
| Methyl dodecanoate | 0 | 0 | 0 | 8,50E+00 |
| Methyl myristate | 0 | 0 | 0 | 5,20E+00 |
| Propyl propionate | 0 | 0 | 0 | 2,02E+01 |
| Propyl butyrate | 1,46E+03 | 2,08E+03 | 1,48E+03 | 6,80E+03 |
| Propyl hexanoate | 3,57E+03 | 1,78E+04 | 3,53E+03 | 8,51E+03 |
| Phenylethyl cinnamate | 1,31E+03 | 1,77E+03 | 1,37E+03 | 2,28E+02 |
| Isobutyl cinnamate | 1,69E+02 | 4,26E+03 | 1,60E+02 | 1,77E+03 |
| Methyl 2,5-dihydroxycinnamate | 0 | 1,37E+01 | 1,02E+01 | 3,06E+01 |
| Methyl cinnamate | 2,28E+02 | 4,04E+02 | 2,34E+02 | 3,33E+03 |
| Methyl ferulate | 1,08E+02 | 2,15E+02 | 1,13E+02 | 5,22E+02 |
| Vinyl acetate | 9,80E+00 | 3,25E+01 | 1,06E+01 | 3,71E+01 |
| Vinyl propionate | 4,60E+00 | 3,25E+01 | 5,10E+00 | 3,25E+01 |
| Vinyl butyrate | 0 | 1,43E+01 | 0 | 3,65E+01 |
| Vinyl laurate | 0 | 5,90E+00 | 0 | 2,02E+01 |
| Vinyl benzoate | 2,06E+03 | 4,56E+03 | 2,07E+03 | 2,75E+04 |
| Vinyl crotonate | 6,31E+02 | 2,15E+03 | 6,87E+02 | 4,78E+03 |
| Vinyl acrylate | 1,54E+03 | 2,82E+03 | 1,54E+03 | 1,30E+03 |
| Geranyl acetate | 1,82E+03 | 7,52E+03 | 1,81E+03 | 1,55E+04 |
| 3-Methyl-3-buten-1-yl acetate | 1,53E+03 | 1,80E+04 | 1,52E+03 | 2,65E+04 |
| Ethyl 2-ethylacetoacetate | 3,12E+01 | 1,30E+01 | 3,57E+01 | 1,83E+02 |
| Ethyl 2-methylacetoacetate | 2,08E+02 | 6,18E+02 | 2,07E+02 | 7,04E+03 |
| Ethyl 3-oxohexanoate | 3,10E+03 | 1,49E+04 | 3,18E+03 | 2,74E+04 |
| Ethyl acetoacetate | 6,83E+02 | 1,93E+03 | 6,89E+02 | 2,88E+03 |
| Ethyl propionylacetate | 7,81E+02 | 2,28E+03 | 7,78E+02 | 1,66E+04 |
| ϒ-Valerolactone | 2,73E+02 | 6,12E+02 | 2,88E+02 | 1,46E+03 |
| Methyl glycolate | 1,39E+02 | 6,38E+02 | 1,38E+02 | 1,35E+04 |
| D-Pantolactone | 1,30E+01 | 2,60E+01 | 1,39E+01 | 3,91E+01 |
| (1*R*)-(-)-dimenthyl succinate | 0,00E+00 | 4,60E+00 | 0,00E+00 | 0 |
| Ethyl 2-chlorobenzoate | 0,00E+00 | 3,12E+01 | 1,44E+01 | 1,56E+02 |
| Cyclohexyl butyrate | 2,96E+03 | 6,50E+03 | 2,96E+03 | 1,98E+04 |
| n-Pentyl benzoate | 5,92E+01 | 9,50E+01 | 6,01E+01 | 2,34E+02 |
| 2,4-Dichlorophenyl 2,4-dichlorobenzoate | 0 | 0 | 0 | 1,24E+01 |
| 2,4-Dichlorobenzyl 2,4-dichlorobenzoate | 2,21E+01 | 1,31E+02 | 2,31E+01 | 1,53E+03 |
| Diethyl-2,6-dimethyl 4-phenyl-1,4-dihydro pyridine-3,5-dicarboxylate | 1,63E+01 | 4,75E+01 | 1,73E+01 | 1,37E+03 |
| (+)-Methyl D-Lactate | 1,30E+00 | 5,21E+01 | 1,50E+00 | 2,35E+04 |
| (-)-Methyl L-Lactate | 2,80E+02 | 1,82E+02 | 2,82E+02 | 2,55E+03 |
| Propyl acetate | 3,90E+00 | 4,56E+01 | 3,90E+00 | 6,20E+03 |
| Butyl acetate | 1,05E+03 | 1,07E+04 | 1,09E+03 | 2,55E+04 |
| Phenyl acetate | 2,59E+04 | 3,12E+04 | 2,63E+04 | 3,26E+04 |
| Phenyl propionate | 3,02E+04 | 2,92E+04 | 3,03E+04 | 3,28E+04 |
| Glucose pentaacetate | 1,69E+02 | 2,60E+02 | 1,73E+02 | 9,50E+02 |

[a]The results are mean from three independent experiments, with SD less than 5% in all cases.

Table S2. Effect of organic solvents on the activity of $EH_0$.

| Solvent | %, (v/v) | Relative activity (%)[b] |
|---|---|---|
| Control[a] | - | 100.0 ± 7.3 |
| Ethanol | 10 | 60.7 ± 9.5 |
| | 20 | 22.3 ± 4.9 |
| | 40 | 1.2 ± 0.4 |
| Methanol | 10 | 159.9 ± 2.9 |
| | 20 | 62.0 ± 9.6 |
| | 40 | 10.6 ± 2.9 |
| Isopropanol | 10 | 20.2 ± 0.6 |
| | 20 | 2.2 ± 0.4 |
| | 40 | 0.6 ± 1.4 |
| Acetonitrile | 10 | 58.9 ± 4.3 |
| | 20 | 18.9 ± 1.2 |
| | 40 | 0.1 ± 0.4 |
| DMSO | 10 | 138.9 ± 5.3 |
| | 20 | 121.9 ± 1.6 |
| | 40 | 15.9 ± 1.8 |
| Acetonitrile/DMSO (50% each) | 1 | 95.1 ± 6.9 |
| | 5 | 99.9 ± 12.6 |
| | 10 | 100.0 ± 6.3 |
| | 20 | 72.4 ± 5.1 |
| | 30 | 32.0 ± 1.4 |
| | 40 | 4.4 ± 0.6 |
| | 50 | 1.3 ± 1.0 |

[a]Reference condition set as 100% activity.

[b]The results are the mean ± SD from three independent experiments.

Table S3. Effect of metal ions of the $EH_0$ esterase. The reactions were performed under standard conditions with the addition of 1, 5, and 10 mM metal salts. The results are expressed as the relative activity of the control sample.

| Cation | Concentration (mM) | Relative activity (%)[b] |
|---|---|---|
| Control[a] | - | 100 |
| $CaCl_2$ | 1 | 98.2 ± 2.5 |
| | 5 | 81.0 ± 8.4 |
| | 10 | 61.7 ± 5.5 |
| $FeCl_3$ | 1 | 89.2 ± 6.7 |
| | 5 | 64.1 ± 7.4 |
| | 10 | 6.8 ± 5.5 |
| $MnCl_2$ | 1 | 94.1 ± 15.5 |
| | 5 | 86.9 ± 2.5 |
| | 10 | 68.9 ± 3.7 |
| $ZnSO_4$ | 1 | 70.6 ± 2.3 |
| | 5 | 69.0 ± 6.8 |
| | 10 | 28.0 ± 6.7 |
| $CuSO_4$ | 1 | 79.2 ± 4.7 |
| | 5 | 57.9 ± 5.0 |
| | 10 | 15.4 ± 0.4 |
| $CoCl_2$ | 1 | 99.9 ± 22.3 |
| | 5 | 81.0 ± 12.0 |
| | 10 | 36.0 ± 0.9 |
| $MgCl_2$ | 1 | 85.0 ± 3.4 |
| | 5 | 101.8 ± 6.6 |
| | 10 | 96.8 ± 4.9 |

[a]The activity of the enzyme without the addition of metal ions was defined as 100%.

[b]The results are the mean ± SD from three independent experiments.

Table S4. Crystallographic statistics of $EH_0$.

| Values in brackets are for the high-resolution shell | |
|---|---|
| **Crystal data** | $\mathbf{EH_0}$ |
| Space group | $P2_12_12_1$ |
| Unit cell parameters | |
| a (Å) | 63.96 |
| b (Å) | 116.67 |
| c (Å) | 128.56 |
| **Data collection** | |
| Beamline | XALOC(ALBA) |
| Temperature (K) | 100 |
| Wavelength (Å) | 1.072180 |
| Resolution (Å) | 64.28-2.01 |
| **Data processing** | |
| Total reflections | 673324 (47503) |
| Unique reflections | 64849 (4496) |
| Multiplicity | 10.4 (10.6) |
| Completeness (%) | 100.0 (100.0) |
| Mean $I/\sigma$ ($I$) | 16.6 (3.6) |
| $R_{merge}^{\dagger}$ (%) | 7.5 (67.7) |
| $R_{pim}^{\dagger\dagger}$ (%) | 2.4 (21.9) |
| Molecules per ASU | 2 |
| **Refinement** | |
| $R_{work}/R_{free}^{\dagger\dagger\dagger}$ (%) | 17.2/20.2 |
| **N° of atoms/average B** (Å$^2$) | 5252/45.92 |
| Macromolecule | 4779/45.57 |
| Ligands | 40/62.44 |
| Solvent | 433/48.26 |
| **Ramachandran plot** (%) | |
| Favored | 95.6 |
| Outliers | 0.9 |
| **RMS deviations** | |
| Bonds (Å) | 0.010 |
| Angles (°) | 1.484 |
| PDB accession code | 7ZR3 |

$^{\dagger}$Rmerge = ∑hkl ∑I | Ii(hkl) – [I(hkl)]|/∑hkl ∑I Ii(hkl), where Ii(hkl) is the ith measurement of reflection hkl and [I(hkl)] is the weighted mean of all measurements.

$^{\dagger\dagger}$Rpim = ∑hkl [1/(N – 1)] ½ ∑I | Ii(hkl) – [I(hkl)]|/∑hkl ∑I Ii(hkl), where N is the redundancy for the hkl reflection.

[†††]Rwork/Rfree = ∑hkl | Fo – Fc |/∑hkl | Fo |, where Fc is the calculated and Fo is the observed structure factor amplitude of reflection hkl for the working/free (5%) set, respectively.

**TABLE S5.** Interface atomic interactions of $EH_0$.

| Hydrogen bonds | | | Salt bridges | | |
|---|---|---|---|---|---|
| **Molecule A** | **Dist. (Å)** | **Molecule B** | **Molecule A** | **Dist. (Å)** | **Molecule B** |
| Leu268(O) | 3.24 | Arg260(NH1) | Glu312(OE1) | 2.87 | Arg276(NH1) |
| Phe275(O) | 3.13 | Val273(N) | Glu312(OE2) | 2.90 | Arg276(NH2) |
| Val273(O) | 2.80 | Phe275(O) | Arg276(NH2) | 2.85 | Glu312(OE2) |
| Arg260(NH1) | 3.05 | Leu268(O) | | | |
| Val273(N) | 3.09 | Phe275(O) | | | |
| Phe275(N) | 2.81 | Val273(O) | | | |
| Lys279(NZ) | 2.70 | Gly270(O) | | | |

**TABLE S6.** Raw data corresponding to calculations, using GraphPad Prism software (version 6.0), of kinetic parameters for p-NP acetate, p-NP butyrate and p-NP dodecanoate. Because its extensive size, the table is provided in a separate Excel table. For kinetic determinations, p-NP esters that are common substrates that denote high robustness, sensitivity, precision and reproducibility on the determinations were used, at concentrations above the detection limits of the available equipment for p-NP (i.e. µM range).

**TABLE S7.** (A-B) Follow up hydrolysis, by the soluble enzymes (wild type and mutants), of a series of 96 carboxylic ester substrates (see panel C) by means of a high-throughput pH indicator assay in 384-well plates. The acid produced after ester bond cleavage by the hydrolytic enzyme induced a color change in the pH indicator that can be measured spectrophotometrically at 550 nm. The table provides the kinetics of the hydrolysis per each ester are shown, followed by recordings of the absorbance at 550 nm over time. Reaction conditions were as described in Materials and Method section, with raw data in the table referred to the tests performed using as stock solution of 0.1 mg/ml enzyme (final enzyme amount in each well, 0.2 µg) (Panel A), or 1.0 mg/ml enzyme (final enzyme amount in each well, 2 µg) (Panel B). Activity (unit/mg protein) is calculated by determining the absorbance per minute from the slopes generated, in the linear range, and by considering an extinction coefficient of phenol red at 550 nm and pH 8.0 of 8450 $M^{-1}cm^{-1}$, a total volume of 44 µL, and a total protein amount of 0.2 µg. Note: as example, the figure representing the kinetic for the first ester is shown in panel A, in which the decrease of absorbance is observed until all ester is consumed and a lag phase is observed (top figure); activity is calculated only considering the linear range shown in the bottom figure. Absence of activity was defined as at least two-fold background signal, with precision in the calculation of two decimal digits [references S1-S3]).

Only the raw data for one of the triplicates are shown. (C) Identity of esters tested in this study. The positions in the 384-well plates are shown in columns A-D. Note that, 96 different esters were tested in each plate, so that 4 tests could be performed (one corresponding to columns 1-6, second columns 7-12, third columns 1318, and fourth columns 19-24), as shown in figure at the bottom of the table (an example of the follow up kinetics (absorbance vs time) is on the right side). Because its extensive size, the table is provided in a separate Excel table.

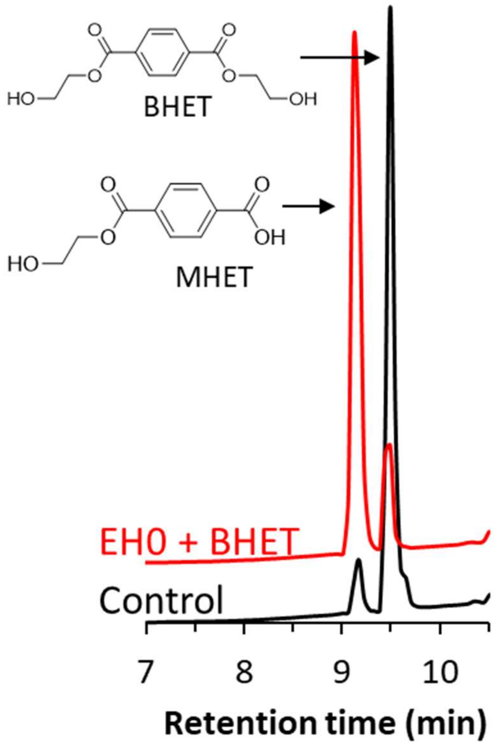

FIG S1. Representative HPLC chromatogram representing the hydrolysis of BHET by $EH_0$. The standards (TA, BHET) and purified (MHET and BHET dimer) products. The figure was made using Excel 2019. High-performance liquid chromatography datasets were collected with a Varian Star LC workstation 6.41 and analyzed using Excel 2019.

```
EH0                 MTELFVRPDVRGFLDFLNNLPGPKMHELDAPTARQMYVAMKDVGDPPVGELGTLLDLSIP
WP_066587239.1      MTEPFVRPDVRGFLDFLNNLPGPKMHELDAPTARQMYVAMKDVGDPPVGELGTLLDLSIP
                    *** ********************************************************

EH0                 GPGGDIPARLYDPRASREPGPAIVFFHGGGFVIGDLESHGSFTAEMARVLDLPVIAVDYR
WP_066587239.1      GPGGNIPARLYDPRASREPGPAIVFFHGGGFVIGDLESHGSFTAEMARVLDLPVIAVDYR
                    ****:*******************************************************

EH0                 LAPEFPWPAAPDDCEAAARWVANSPAELGRSVTSLVLCGDSAGGNLVIVTAAALRDQPAK
WP_066587239.1      LAPEFPWPAAPDDCEAAARWVANSPVELGRSVTSLVLCGDSAGGNLVIVTAAALRDQPAK
                    *************************.**********************************

EH0                 VPVIAQLPFYPATDASKEYPSYAEFAEGYLLTRDSMEWFMAAYKSEADHIRSSPLLGDLA
WP_066587239.1      VPVIAQLPFYPATDASKEYPSYAEFAEGYLLTRDSMEWFMAAYKSEADHIRSSPLLGDLA
                    ************************************************************

EH0                 GMPPAVVVTGGLDPIRDQGRAYAAALALAGVPVVFREAKGNIHGFITLRKAIPSSVGDVM
WP_066587239.1      GMPPAVVVTGGLDPIRDQGRAYAAALALAGVPVVFREAKGNIHGFITLRKAIPSSVGDVM
                    ************************************************************

EH0                 GAFAALKDIIVEAEGDRAMAQAAA
WP_066587239.1      GAFAALKDIIVEAEGDRAMAQAAA
                    ************************
```

FIG S2. Alignment of WP_066587239 and $EH_0$ sequences, using Clustal 2.1 Omega sequence alignment program.

### References

S1. Martínez-Martínez M, Coscolín C, Santiago G, Chow J, Stogios PJ, Bargiela R, Gertler C, Navarro-Fernández J, Bollinger A, Thies S, Méndez-García C, Popovic A, Brown G, Chernikova TN, García-Moyano A, Bjerga GEK, Pérez-García P, Hai T, Del Pozo M V., Stokke R, Steen IH, Cui H, Xu X, Nocek BP, Alcaide M, Distaso M, Mesa V, Peláez AI, Sánchez J, Buchholz PCF, Pleiss J, Fernández-Guerra A, Glöckner FO, Golyshina O V., Yakimov MM, Savchenko A, Jaeger KE, Yakunin AF, Streit WR, Golyshin PN, Guallar V, Ferrer M. 2018. Determinants and prediction of esterase substrate promiscuity patterns. ACS Chem Biol 13:225–234.

S2. Giunta CI, Cea-Rama I, Alonso S, Briand ML, Bargiela R, Coscolín C, Corvini PFX, Ferrer M, Sanz-Aparicio J, Shahgaldian P. 2020. Tuning the properties of natural promiscuous enzymes by engineering their nano-environment. ACS Nano 14:17652–17664.

S3. Nutschel C, Coscolín C, David B, Mulnaes D, Ferrer M, Jaeger KE, Gohlke H. 2021. Promiscuous esterases counterintuitively are less flexible than specific ones. J Chem Inf Model 61:2383-2395.
